# Fungi activate Toll-1 dependent immune evasion to induce cell loss in the host brain

**DOI:** 10.1101/2024.04.29.591341

**Authors:** Deepanshu N.D. Singh, Abigail R.E. Roberts, Enrique Quesada Moraga, David Alliband, Elizabeth Ballou, Hung-Ji Tsai, Alicia Hidalgo

**Affiliations:** Brain plasticity & Regeneration Lab, School of Biosciences, University of Birmingham, UK; Institute of Immunity and Infection, School of Biosciences, University of Birmingham, UK; Current address: Soller Lab, School of Biosciences, University of Birmingham, UK; Current address: School of Biomedical Sciences, University of Leeds, UK; Current address: University of Bristol, UK; Departamento de Agronomía, Universidad de Córdoba, Spain; Current address: MRC Centre for Medical Mycology, Geoffrey Pope Building, University of Exeter

**Keywords:** *Drosophila*, Beauveria bassiana, fungal infection, Toll-1, spz, wek, sarm, host-pathogen interactions, neurodegeneration, dopamine, dopaminergic neuron

## Abstract

Fungi evolve within the host, ensuring their own nutrition and reproduction, at the expense of host health. They intervene in hosts’ brain function, to alter host behaviour and induce neurodegeneration. In humans, fungal infections are emerging as drivers of neuroinflammation, neurodegenerative diseases and psychiatric disorders. However, how fungi alter the host brain is unknown. Fungi trigger an innate immune response mediated by the Toll-1/TLR receptor, the adaptor MyD88 and the transcription factor Dif/NFκB, that induce the expression of antimicrobial peptides (AMPs). However, in the nervous system, Toll-1/TLR could also drive an alternative pathway involving the adaptor Sarm, which causes cell death instead. Sarm is the universal inhibitor of MyD88 and could drive immune evasion. The entomopathogenic fungus *Beauveria bassiana* is well-known to activate Toll-1 signalling in innate immunity in *Drosophila*. In fruit-flies, the adaptor Wek links Toll-1 to Sarm. Thus, here we asked whether *B. bassiana* could damage the *Drosophila* brain via Toll-1, Wek and Sarm. We show that exposure to *B. bassiana* reduced fly lifespan and impaired locomotion. *B. bassiana* entered the brain and induced the up-regulation of *AMPs,* as well as *wek* and *sarm,* within the brain. Exposure to *B. bassiana* caused neuronal and glial loss in the adult *Drosophila* brain. Importantly, RNAi knockdown of *Toll-1, wek* or *sarm* concomitantly with infection prevented *B. bassiana* induced cell loss. By contrast, over-expression of *wek* or *sarm* was sufficient to cause dopaminergic neuron loss in the absence of infection. These data show that *B. bassiana* caused cell loss in the host brain via Toll-1/Wek/Sarm signalling driving immune evasion. We conclude that pathogens can benefit from an innate immunity receptor to damage the host brain. A similar activation of Sarm downstream of TLRs in response to fungal infections could underlie psychiatric and neurodegenerative diseases in humans.

## INTRODUCTION

Fungi such as *Ophiocordyceps unilateralis* and *Entomopthora muscae* can manipulate the behaviour of their insect hosts, ultimately causing their death (de Bekker, 2019, de Bekker and Das, 2022, Elya et al., 2018). Even only exposure to fungal volatiles is sufficient to reduce lifespan, decrease dopamine levels, induce neurodegeneration and impair locomotion in fruit-flies (Inamdar et al., 2013, Inamdar et al., 2010). This is concerning also for human health. Abundant fungal spores, mould and fungal volatiles are commonly found in indoor damp conditions (Provost et al., 2013, Shi et al., 2014). Fungal spores have been found in brains of Parkinson’s and Alzheimer’s disease patients, and fungal infections are emerging as drivers of neuroinflammation, neurodegenerative diseases and psychiatric disorders (Chen et al., 2022, Alonso et al., 2014, Pisa et al., 2020, Cannon and Gruenheid, 2022). How fungi and neuroinflammation drive disease in the host brain is unknown (Cannon and Gruenheid, 2022, Chen et al., 2022).

In the fruit-fly *Drosophila*, defence against fungi is controlled by the innate immunity Toll signalling pathway (Imler and Hoffmann, 2001, Lemaitre et al., 1996, Leulier and Lemaitre, 2008). Of the nine different *Toll* receptors, Toll-1 has the most prominent function in innate immunity (Tauszig et al., 2000, Chowdhury et al., 2019, Lemaitre et al., 1996). In mammals, homologues of Tolls are the Toll-Like Receptors (TLRs), and Tolls and TLRs across organisms have a universally conserved function in innate immunity (Leclerc and Reichhart, 2004, Leulier and Lemaitre, 2008, Hoffmann and Reichhart, 2002, Gay and Gangloff, 2007b). Mammalian TLRs are pattern recognition receptors (PRR) that directly bind pathogens (Gay and Gangloff, 2007a), but in Drosophila fungi are recognised by GNBP3 instead (Gottar et al., 2006). This initiates a proteolytic cascade that activates Spätzle Processing Enzyme (SPE) that cleaves Spätzle (Spz), ligand of Toll-1 (Gay and Gangloff, 2007a, Hu et al., 2004, Jang et al., 2006, Weber et al., 2003, Matskevich et al., 2010). Toll-1 signalling proceeds via MyD88, triggering the nuclear translocation of the transcription factor Dif/NF-κB (Anthoney et al., 2018, Ip et al., 1993, Lemaitre et al., 1996, Imler and Hoffmann, 2001). Dif activates the expression of antimicrobial peptide (AMP) encoding genes, such as *drosomycin (drs)* and *metchnikowin (mtk),* that protect *Drosophila* from fungal infections (Brennan and Anderson, 2004, Fehlbaum et al., 1994, Hanson et al., 2021, Lemaitre et al., 1996, Zhang and Zhu, 2009, Volkoff et al., 2003).

At least seven of the nine *Toll* receptor paralogues are expressed in the adult fly brain (McIlroy et al., 2013, Foldi et al., 2017, Li et al., 2020). In the nervous system, Tolls have functions unrelated to immunity, in axonal connectivity, neurogenesis, cell survival, cell death, structural brain and synaptic plasticity (Li et al., 2020, Ward et al., 2015, Foldi et al., 2017, McIlroy et al., 2013, McLaughlin et al., 2016, Ballard et al., 2014). Not all Tolls function equally, and Toll-1 is more likely to drive apoptosis than others (McIlroy et al., 2013, Foldi et al., 2017, Li et al., 2020). Toll receptors can either promote cell survival via Wek-MyD88-NFκB or cell death via Wek-Sarm-JNK signalling downstream (Foldi et al., 2017, McIlroy et al., 2013, Li et al., 2020). Sarm is the universal, highly conserved, only inhibitor of MyD88 and TRIF signalling (Anthoney et al., 2018, Belinda et al., 2008, Carty and Bowie, 2019, Peng et al., 2010, Yuan et al., 2010). It can be at the plasma membrane and associated with mitochondria, it can induce neuronal death via inhibiting the pro-survival function of MyD88, via activating JNK signalling and via NADase catalytic activity, which also drives neurite self-destruction (Anthoney et al., 2018, Carty et al., 2006, Coleman and Freeman, 2010, Conforti et al., 2014, Essuman et al., 2017, Foldi et al., 2017, Izadifar et al., 2021, Osterloh et al., 2012, Sarkar et al., 2023, Mukherjee et al., 2015). In the brain, signalling from Toll receptors can activate three distinct signalling pathways that result in four alternative cellular outcomes – cell survival, cell death, cell quiescence or cell proliferation – depending on cell context (Foldi et al., 2017, Li et al., 2020).

Here, we asked whether fungal infections could induce neurodegeneration in the host brain via Toll-Wek-Sarm signalling. We used *Beauveria bassiana,* an entomopathogenic fungus that induces disease in over 700 arthropod species, can be used for the control of insect pests, and is a well-known activator of Toll signalling in *Drosophila* (Evans, 1982, Gottar et al., 2006, Inglis et al., 2001, Lemaitre et al., 1997, McCoy, 1990, Samson et al., 1988, Quesada-Moraga et al., 2023). We show that the fungus benefits from immune receptor signaling as it induces immune evasion and cell death in the host brain.

## RESULTS

### *B. bassiana* exposure decreased lifespan and impaired locomotion

To ask whether and how fungi could affect the brain, we aimed to mimic natural exposure of flies to spores. We used an infection chamber with a fungal culture at the base and a separate compartment for fly-food affixed to the bottle’s side wall, thus preventing fungal growth from limiting food and hydration supply (Figure 1A). Whereas non-infected control flies lived up to 70 days, flies exposed to *B. bassiana* died within less than 20 days (Figure 1B). By day seven over 50 percent of the flies in the infection chamber had perished (Figure 1B). Next, we tested whether exposure to *B. bassiana* affected fly behaviour. Negative geotaxis-also known as startle induced negative geotaxis (SING) or the climbing assay-is commonly used to measure locomotor impairment caused by neurodegeneration (Feany and Bender, 2000, Riemensperger et al., 2013). Amongst the surviving flies, three days exposure to *B. bassiana* had no effect on flies’ climbing ability, but by seven days post exposure, climbing was significantly impaired (Figure 1C). Thus, from here onwards, we used three-and seven-days’ exposures for further tests.

**Figure 1.**
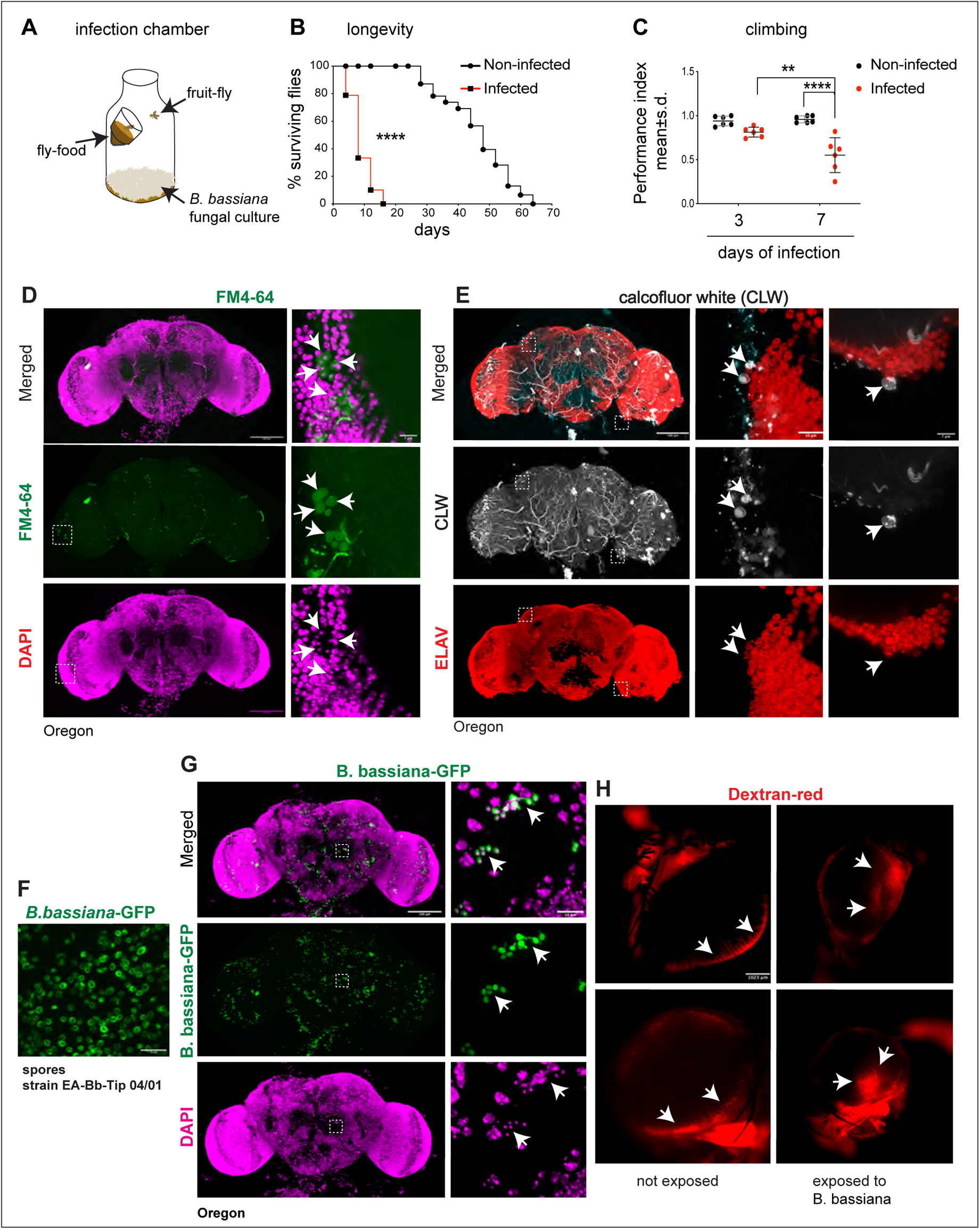
*B. bassiana* spores were detected within the fly brain. **(A)** Infection chamber. **(B)** *B. bassiana* exposure reduced lifespan. Percentage surviving flies at each scored day Log-rank test p<0.0001 n=104 non-infected flies, and infected flies n=104. **(C)** *B. bassiana* exposure impaired climbing. Dot plots, with lines indicating the mean and standard deviation. Each dot represents one cohort of 7-10 flies. Two-way ANOVA: 3-days-old vs 7-days-old p= 0.0122; Noninfected vs infected: p<0.0001; Interaction: p= 0.0048. 3 days old: n= 60 for each infected and non-infected. 7 days old: n= 51 for both infected and non-infected. Turkey’s multiple comparison correction test: **p< 0.01, ***p<0.001. **(D,E,G)** *B. bassiana* infiltrates the *Drosophila* adult brain as visualised with: **(D)** FM4-64 dye, revealing co-localisation with DAPI+ nuclei in spores. **(E)** Calcofluor white, revealing spores adjacent to neurons within the central brain and optic lobes. **(F)** A culture of *B. bassiana*-GFP strain EA-Bb-Tip 04/01. **(G)** Transgenic *B. bassiana* GFP, revealing GFP+ spores within the central brain. **(H)** *B. bassiana* damaged the blood brain barrier in the eye, as dextran red could diffuse within the retina in infected brains (arrows) but not in non-infected controls. Scale bars: (C, D, E) left: 100μm; (C,D) right: 7μm; (D) middle, (E) right and (F): 10μm; (G) 1023 μm. Sample sizes: non-infected n = 6, Infected n = 5

Altogether, these data showed that exposure to *B. bassiana* compromised locomotion and longevity, which are common indicators of neurodegeneration in flies.

### *B. bassiana* penetrate adult fly brains

In the light of the locomotion impairment, we wondered whether *B. bassiana* could affect brain function directly, by entering the brain. We exposed adult flies to *B. bassiana* spores for three days and using the FM4-64 dye, distinctive oval-shaped structures could be seen within the brain optic lobes (Figure 1D). Importantly, here, FM4-64 colocalised with the nuclear dye DAPI, confirming that they were spores (Figure 1D). We used a second dye, calcofluor white (CLW), which stains cell walls of algae, plants and fungi, together with the pan-neuronal marker Elav, and CLW+ signal was found in the optic lobes and central brain after three days of *B. bassiana* exposure (Figure 1E). Finally, when adult flies were exposed to a GFP transgenic *B. bassiana* strain (EABb 04/01-Tip GFP5, Figure 1F) (Landa et al., 2013), GFP+ spores were found both within the optic lobes and central brain (Figure 1G). Together these findings show that *B. bassiana* spores infiltrated the adult brain.

The blood brain barrier (BBB) shelters the brain from bodily fluids and foreign agents, thus, to enter the brain, *B. bassiana* would either have to damage the BBB or be carried across by macrophages (Winkler et al., 2021). The dye dextran red is commonly used to test the integrity of the BBB in *Drosophila* (Bainton et al., 2005). The dye is injected into the flies’ thorax, and if the BBB is intact, dextran red outlines the retina’s edge, whereas if the integrity of the BBB is compromised, the dye leaks into the retina (Bainton et al., 2005). We found that in non-exposed flies the dye accumulated in the retina’s periphery, whereas in flies that had been exposed to *B. bassiana* for seven days, dextran red had spread within the retina (Figure 1H), meaning that the BBB was damaged.

Altogether, these findings showed that following exposure, *B. bassiana* penetrated the fruit-fly brain.

### Exposure to *B. bassiana* activated Toll signalling inside the adult brain

*B. bassiana* could invade the brain through the optic lobes, the head capsule, and the proboscis. Fungi can degrade insects’ cuticle, enabling them to penetrate hosts’ bodies (Boucias et al., 1988, Holder et al., 2007, Ortiz-Urquiza and Keyhani, 2016, Quesada-Moraga et al., 2020). The fly uses the proboscis for feeding, and as it traverses the brain to join the oesophagus, *B. bassiana* could enter the brain through the proboscis during feeding. On the other hand, conceivably, *B. bassiana* could trigger repulsion or avoidance in the fly. In *C.elegans* worms, pathogenic bacteria trigger a behavioural avoidance response, inducing the worms to turn away from the source of pathogen (Pradel et al., 2007, Pujol et al., 2001). Thus, we tested whether flies like or dislike to feed on *B. bassiana* spores. We offered them either water or sucrose or spores mixed with a blue dye and measured the optical density (OD) of their abdomens after feeding. OD values were higher in flies that fed on spores compared to those that fed on water controls. This showed that flies fed on sucrose – which they are known to like – and similarly fed abundantly on spores (Figure 2A). Thus, ingestion would provide *B. bassiana* with an effective route into the brain.

**Figure 2.**
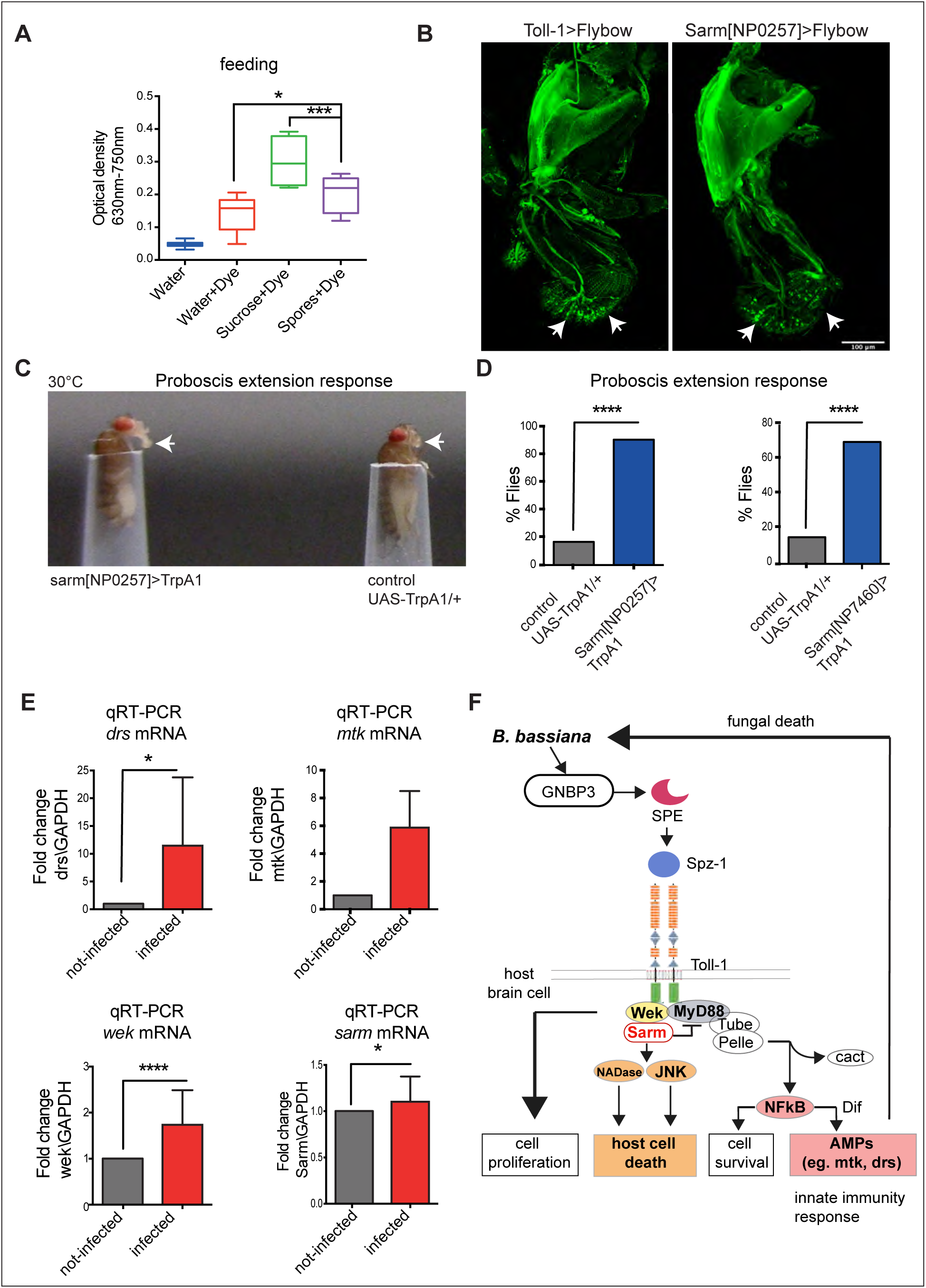
Exposure to *B. bassiana* activated Toll signalling within the adult brain. **(A)** *Drosophila* ate *B. bassiana* spores that had been mixed with blue dye diluted in water. One Way ANOVA P value < 0.0001. **(B)** *Toll-1>FlyBow* and *Sarm^NP0257^>FlyBow* reveal expression in the proboscis, note axons and dendrites of sensory neurons in the labellum (arrows). n=3-7 brains. **(C)** Activating Sarm+ neurons triggered the proboscis extension response, a proxy for feeding. Snapshot of films. **(D)** Quantification of **(C)** in bar charts. Chi-square test, ****p<0.0001. Test: Sarm^NP0257^>TrpA1 n=34 flies, Control: +(Oregon)>TrpA1 n=34 Test: Sarm^NP7460^>TrpA1 n=40, Control: +(oregon)>TrpA1 n=40. **(E)** *B. bassiana* infection for 7 days raised the expression levels of AMPs within the brain. qRT-PCR data comparing fold change levels of *drs* and *Mtk* mRNA (top) and wek and sarm (bottom) in infected vs. non-infected brains, normalized to GAPDH as a housekeeping control. DeltaCT mean ± standard deviation. *drs:* Unpaired student t-test, *p<0.05, n=3 biological replicates (b.r.); mtk: p=0.0634; n=3 b.r.; *wek:* Unpaired student t-test ****p<0.0001 n=4 b.r.; *sarm:* t test: *p<0.05, n = 4 b.r.. **(F)** Toll-1 signalling drives three distinct signalling pathways downstream, that result in different cellular outcomes. Sarm also inhibits MyD88 signalling. Scale bar in (B): 100 μm

Above findings showed *B. bassiana* inside the brain, that flies were not repelled by *B. bassiana* and instead fed on it. This was surprising for two reasons. Firstly, it contrasts with the bacterial avoidance response of *C.elegans* which depends on Tol-1 (Pradel et al., 2007, Pujol et al., 2001). Secondly, in *Drosophila*, the innate immune response driven by Toll-1 should lead to clearance of the fungus (Imler and Hoffmann, 2001). Thus, we wondered whether perhaps *B. bassiana* did not lead to the activation of Toll-1 inside the brain. First, we asked whether *Toll-1* and *sarm* might be expressed in the proboscis, which is required for feeding. If they were not, this would explain the lack of avoidance response to *B. bassiana*. We had previously shown that both *Toll-1* and *sarm* are expressed in the sub-aesophageal ganglion, where the feeding circuit links to the proboscis (Li et al., 2020). Here, using the in vivo reporter *Toll-1>FlyBow* we found expression in sensory neurons of the proboscis, including in dendrites of the labellum (Figure 2B). Similarly, the Toll signalling adaptor *sarm* was also expressed in sensory neurons of the proboscis (Figure 2B). To further verify this, we tested what consequences activating Sarm+ neurons might have on the proboscis extension response (PER), which is required for feeding. Activating Sarm neurons with TrpA1 (using two independent lines) did not prevent feeding, and instead increased the incidence of PER events compared to unstimulated controls (Figure 2C,D). This suggests that *sarm* neurons are not involved in avoidance of *B. bassiana*, but instead could be involved in feeding.

Next, we asked whether *B. bassiana* could trigger Toll signalling in the brain, which would result in the up-regulation of its immunity downstream effectors. We had previously shown that seven of the nine *Toll* receptors, as well as its adaptors *MyD88* and *sarm*, are normally expressed in the adult fly brain (Li et al., 2020). Single-cell RNAseq analysis of the adult fly brain revealed that the fungal PRR *GNBP3* as well as the downstream *SPE* protease, *spz-1* ligand of *Toll-1*, downstream Toll-1 adaptor *MyD88*, downstream effector *Dif/NF-κB* and target antimicrobial peptide genes *drs* and *mtk*, are all normally expressed in the adult brain (Scope Fly Atlas)(Davie et al., 2018). Thus, we asked whether exposure to *B. bassiana* might activate innate immunity in the brain. We used qRT-PCR in dissected adult brains, and found that at seven days post-infection, expression of AMPs *drs* and *mtk* mRNA was upregulated (Figure 2E). Intriguingly, the expression of the Toll adaptors *wek* and *sarm* was also upregulated (Figure 2E). Similarly, *wek* expression was also shown to increase upon *E. muscae* infection in adult flies (Elya et al., 2018), consistent with our findings. To conclude, our data showed that exposure to *B. bassiana* induced immune Toll signalling within the adult brain. Moreover, this supported the data shown in (Figure 1) that *B. bassiana* entered the brain. Intriguingly, the data showed that *B. bassiana* also triggered an alternative Toll signalling route engaging Wek and Sarm (Figure 2F). Wek can interact with MyD88, but it is not required for innate immunity (Chen et al., 2006), and it links Tolls to Sarm instead (Foldi et al., 2017). Sarm inhibits MyD88 and immune signalling, and instead, Toll signalling via Wek and Sarm induces apoptosis downstream (Foldi et al., 2017). Sarm also induces axonal destruction (Osterloh et al., 2012). Remarkably, the infection-dependent up-regulation of *wek* and *sarm* had the potential to drive neurodegeneration in response to Toll signalling (Figure 2F).

### Exposure to *B. bassiana* caused neuronal and glial loss

To ask whether *B. bassiana* could induce neurodegeneration via Toll-1, Wek and Sarm, we first tested whether exposure to *B. bassiana* altered cell number in the brain. If Sarm drove neurodegeneration in response to infection, these cells would be lost. Thus, to test if the number of Sarm+ neurons was affected, we used the nuclear reporter histone-YFP to visualize Sarm+ cells in the central brain (*sarm^NP0257^>hisYFP*) and quantified Sarm+ cells automatically. Seven days exposure to *B. bassiana* decreased the number of Sarm+ cells in the central brain compared to non-exposed controls (Figure 3A,B). To ask whether glial cells might also be affected, glial cells were labelled with pan-glial anti-Repo antibodies, and we found that the number of glial cells in infected flies also decreased (Figure 2C,D). Thus, *B. bassiana* infection caused loss of Sarm+ neurons and Repo+ glia.

**Figure 3.**
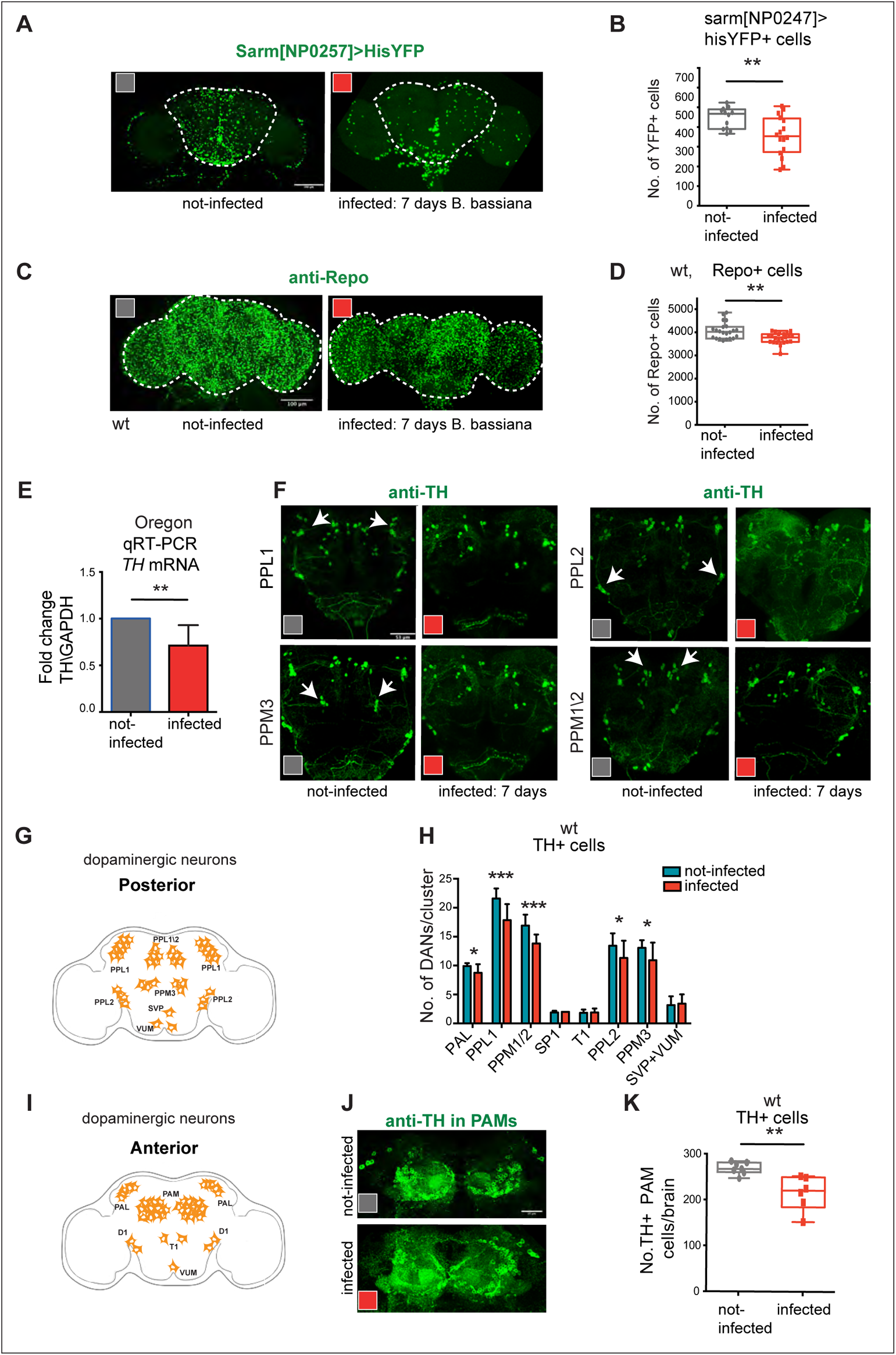
Exposure to *B. bassiana* caused cell loss in the fly brain. **(A,B)** Seven-day exposure to the *B. bassiana* caused loss of *sarm^NP0257^>HisYFP* cells. Quantification in **(B),** Mann-Whitney U test, p=0.002. **(C,D)** Seven-day exposure to the *B. bassiana* caused loss of glia cells, visualised with anti-Repo antibodies. Quantification in **(D),** Unpaired Student t-test, p=0.002. **(E)** Seven-day exposure to the *B. bassiana* reduced tyrosine hydroxylase (TH) expression. qRT-PCR showing fold-change relative to non-infected wild type controls and normalised to GAPDH, mean ± standard deviation. Unpaired Student t-test on delta-Ct values p=0.0011, n=3 biological replicates. **(F)** Seven-day exposure to the *B. bassiana* caused loss of dopaminergic neurons (DANs) visualised with anti-TH antibodies, in the posterior brain, quantification in **(H)**: PAL, SP1, T1: Mann-Whitney U tests; PPL1, PPM1/2, PPL2, PPM3, SVP, VUM: Student t tests; non-infected brains n=12, infected brains n=12**. (G,I)** Illustration of dopaminergic neurons in the adult brain. **(J,K)** Seven-day exposure to the *B. bassiana* caused loss of PAMs, quantification in **(K):** unpaired Student t-test, p=0.0043, non-infected brains n=7, infected brains n=6. Graphs in (B,D,K) show box-plots around the median, box with 50% of data points and whiskers with 25% of data points. Graphs in (E,H) show bar charts with mean ± standard deviation. Dotted lines in (A,C) indicate ROI for automatic cell counting with DeadEasy software. **p<0.01, ****p< 0.0001. Scale bars: (A,C,F): 100μm. (J) 25μm.

Exposure of fruit-flies to the fungal volatile 1-octen-3-ol caused loss of dopaminergic neurons (Inamdar et al., 2013). Thus, we asked whether exposure to *B. bassiana* spores could elicit similar effects. First, we tested whether seven-day exposure to *B. bassiana* spores might affect the expression of Tyrosine Hydroxylase (TH), the enzyme that catalyses the conversion of tyrosine to L-Dopa, which is then decarboxylated to produce dopamine (Daubner et al., 2011). Using qRT-PCR, we found a significant decrease in *TH* mRNA levels within infected adult brains (Figure 3E), showing that *B. bassiana* exposure decreased dopamine production. Next, we used anti-TH antibodies to visualise dopaminergic neurons in the adult brain. By seven days post-exposure, the number of PPL1, PPL2, PPM1/2, PPM3 (Figure 3F-I), and PAM (Figure 3I,J,K) dopaminergic neurons had decreased in infected brains. These data show that exposure to *B. bassiana* caused dopaminergic neuron cell loss within the adult brain.

Altogether, seven days exposure to *B. bassiana* caused loss of Sarm>his-YFP+ neurons, Repo+ glia and TH+ dopaminergic neurons in the infected adult fly brain.

### *B. bassiana* requires Toll-1 to induce cell loss in the fly brain

Having established that exposure to *B. bassiana* induced Toll-1 signalling and caused cell loss in the adult brain, we asked whether cell loss depended on Toll-1. Raised levels of anti-microbial peptides of the IMD pathway, as well as *drs* of the Toll-1 pathway, can induce neurodegeneration in the fly brain (Cao et al., 2013, Kounatidis et al., 2017, Shukla et al., 2019). On the other hand, Toll-1 signalling can induce cell death via the Wek/Sarm/JNK pathway (Foldi et al., 2017). Sarm is the only inhibitor of MyD88, and this function is highly evolutionarily conserved (Belinda et al., 2008, Carty et al., 2006, Peng et al., 2010, Carty and Bowie, 2019). Thus, in MyD88+ cells, Toll-1 could potentially drive signalling via the two alternative MyD88 and Sarm pathways, in co-expressing cells. Thus, to account for both potential Toll-1 signalling routes, we visualised adult MyD88+ cells with *MyD88>HisYFP* and counted cell number automatically. Seven-day exposure to *B. bassiana* induced loss of MyD88+ cells in an otherwise wild-type background (Figure 4A,C).

**Figure 4.**
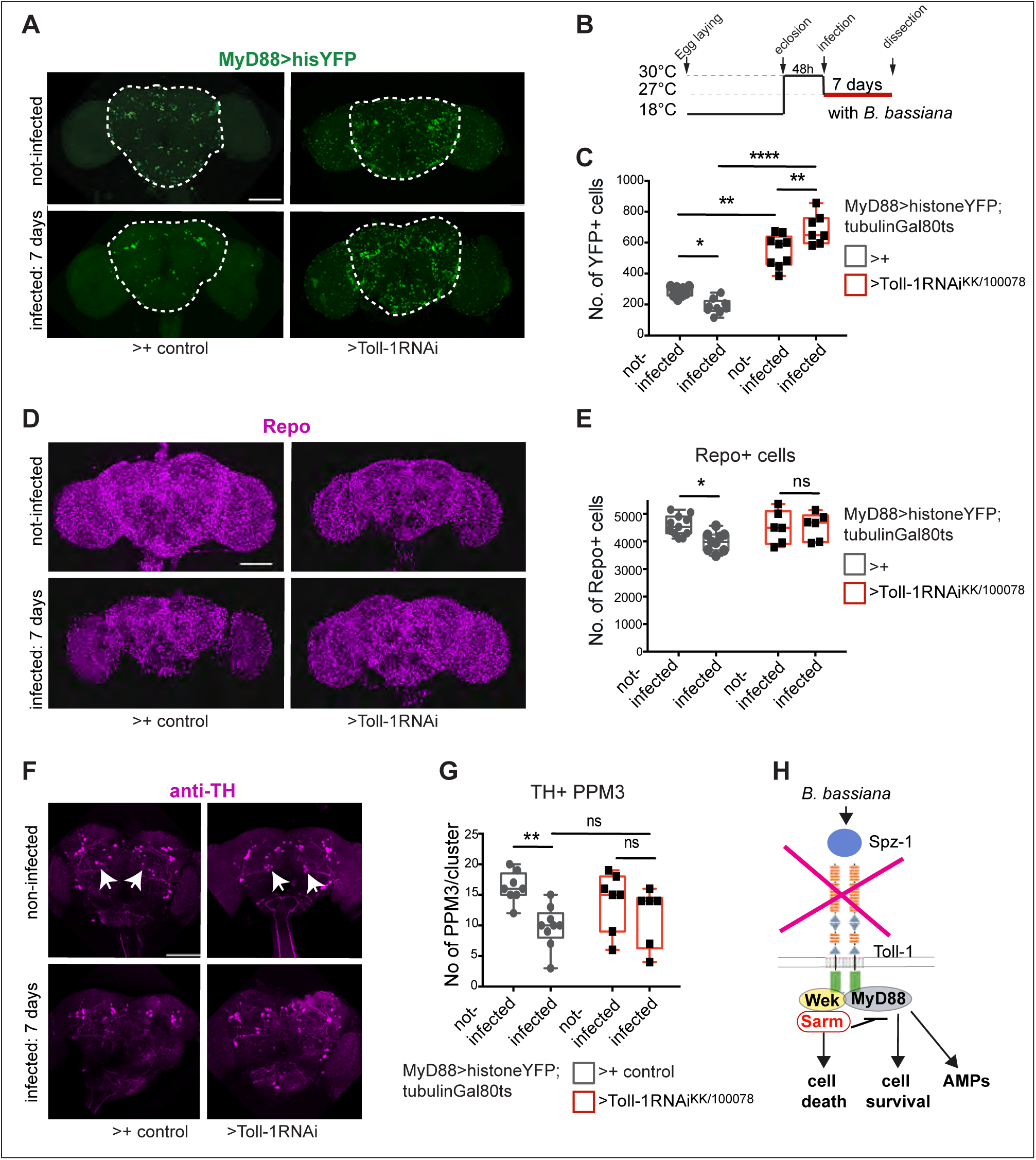
*B. bassiana* induced cell loss depends on Toll-1. **(A)** Seven-day exposure to *B. bassiana* caused loss of MyD88+ cells, visualised with *tubGAL80^ts^; MyD88>HisYFP,* and this was rescued with adult-specific *Toll-1-RNAi* ^KK/100078^ knockdown**. (B)** Diagram explaining the experimental temperature regime. **(C)** Quantification of data in (A). Two-way ANOVA: Infected vs not-infected:p= 0.7557; Control vs Toll-1RNAi (*tubGAL80^ts^; MyD88>HisYFP>Oregon* vs *tubGAL80^ts^; MyD88>HisYFP>Toll-1-RNAi* ^KK/100078^): p<0.0001; Interaction: p<0.0001. Turkey’s multiple comparison correction test. Sample sizes: non-infected control brains n=10, infected control brains n=9, non-infected Toll-1-RNAi brains n=9, infected Toll-1RNAi brains n=9. **(D, E)** Seven-day exposure to *B. bassiana* caused loss of glial cells visualised with anti-Repo antibodies **(D),** which was rescued with adult-specific *Toll-1-RNAi* ^KK/100078^ knockdown, quantification in **(E).** Two-way ANOVA: Infected vs not-infected: p= 0.1003; Control vs Toll-1RNAi p= 0.1285; interaction: p= 0.0666. Turkey’s multiple comparisons correction test. Sample sizes: non-infected control brains n=11, infected control brains n=8, non-infected Toll-1-RNAi brains n=6, infected Toll-1-RNAi brains n=6. **(F,G)** Seven-day exposure to *B. bassiana* caused loss of PPM3 DANs (arrows), visualised with anti-TH antibodies and this phenotype was rescued with adult-specific *Toll-1-RNAi* ^KK/100078^ knockdown. Quantification in **(G).** Two-way ANOVA: Infected vs not-infected: p= 0.0042; Control vs Toll-1 RNAi: p= 0.8548; interaction: p= 0.1760, and Turkey’s multiple comparisons correction test. Sample sizes: non-infected control brains n=8, infected control brains n=9, non-infected Toll-1-RNAi brains n=7, infected Toll-1-RNAi brains n=6. **(H)** Illustration showing how elimination of Toll-1 through adult-specific knock-down prevented cell death outcome after infection. Dotted line in (A) indicates ROI for automatic cell counting with DeadEasy. Graphs in (C,E,G) show box-plots around the median, box with 50% of data points and whiskers with 25% of data points. Asterisks on graphs indicate multiple comparison correction tests: *p<0.05, **p<0.01, ****p< 0.0001. Scale bars: (A, D, F) 100 μm.

To knock-down Toll-1 specifically in adult flies, we used the temperature-sensitive Gal4 repressor tubulin-Gal80ts (Figure 4B, *tubGAL80ts, MyD88>hisYFP*). In non-infected controls, adult *Toll-1* RNAi knockdown using line *UAS-Toll-1RNAi^kk/100078^* (shown to work effectively in (Li et al., 2020)) caused an increase in MyD88-YFP+ cells within the central brain (Figure 4A-C). This increase in cell number could correspond to the induction of proliferation or an increase in the number of surviving cells. In fact, there is abundant cell death in the developing CNS and Toll-1 is known to induce apoptosis in at least larvae and pupae (Foldi et al., 2017). There, Toll-1 loss of function increased neuronal number, and constitutively active *Toll-1^10b^* decreased neuron number and increased the number of Dcp1+ apoptotic cells (Foldi et al., 2017). This suggests that the increase in cell number caused by *Toll-1-RNAi* knock-down would be more likely to result from decreased cell death. Importantly, in infected adult brains, *Toll-1 RNAi* knockdown prevented the loss of MyD88-YFP+ cells that would have been otherwise caused by *B. bassiana* exposure (Figure 4A-C). These data mean that *B. bassiana* induced cell loss depends on Toll-1 signalling.

Neuronal loss can cause glial loss (Hidalgo et al., 2001), *MyD88* is also expressed in some Repo+ glial cells (Li et al., 2020) and *Toll-1* is also expressed in glial cells (Davie et al., 2018). Thus, we tested whether adult specific *Toll-1* RNAi knockdown in MyD88 cells affected glial cell number, stained with anti-Repo. When *Toll-1* was downregulated in non-infected flies, the number of Repo+ glial cells in the brain was not altered compared to controls (Figure 4D,E). However, *Toll-1* RNAi knock-down in MyD88+ cells prevented the decrease in glial cell number caused by *B. bassiana* infection (Figure 4D,E). This meant that *B. bassiana* induced glial cell loss requires Toll-1 signalling in MyD88+ cells.

We also tested the effect of *Toll-1* RNAi knock-down in dopaminergic neurons (DANs). We first tested if *Toll-1* is expressed in DANs, by labelling Toll-1 cells with *Toll-1> HistoneYFP* and dopaminergic neurons with anti-TH antibodies. There was colocalization within multiple DAN clusters, most particularly within the PAL, PPM3, PPL2, and PPL1 clusters (Supplementary Figure 1A). By contrast, very few neurons within the large PAM clusters were Toll-1>hisYFP+ (Supplementary Figure 1A). Following *Toll-1* RNAi knock-down in MyD88+ cells in the adult brain, we counted the number of cells in each DAN cluster. In non-infected flies, *Toll-1* knockdown did not alter DAN cell number (Figure 4F,G). By contrast, *Toll-1* RNAi knockdown prevented infection-induced neuronal loss within the TH+ PPM3 and PPL1 DAN clusters (Figure 4G and Supplementary Figure 1B). This showed that DAN degeneration upon *B. bassiana* infection required Toll-1 (Figure 4H).

### *B. bassiana* benefits from non-immune Wek-Sarm signalling to induce cell loss

Apoptotic Toll-1 signalling requires Wek, which binds Sarm, which induces cell death (Foldi et al., 2017). As we had seen that exposure to *B. bassiana* induced the upregulation of *wek* and *sarm* in the fly brain (Figure 2D), and *B. bassiana* induced cell loss depended on Toll, we asked whether *wek* and *sarm* might also be required for infection-dependent cell loss.

We first asked whether adult specific knock-down of *wek* might affect the cellular response to *B. bassiana* infection. In non-infected controls, adult specific knock-down using line *UAS-wek ^MH046534^-RNAi* (shown to work effectively in (Foldi et al., 2017, Li et al., 2020)) in MyD88+ cells did not alter MyD88-hisYFP+ cell number (Figure 5A,C). However, when *wek* was knocked-down with RNAi, this rescued *B. bassiana*-induced MyD88-HisYFP+ cell loss (Figure 5A,C). In fact, like before, cell number increased further. In non-infected control flies, *wek*-RNAi knockdown did not alter the number of Repo+ glial cells in the brain (Figure 5D,E). However, no difference was found in Repo+ glial cell number between infected and non-infected brains upon *wek*-RNAi knock-down (Figure 5D,E). These data means that *B. bassiana* could not induce MyD88+ and Repo+ cell loss without Wek.

**Figure 5.**
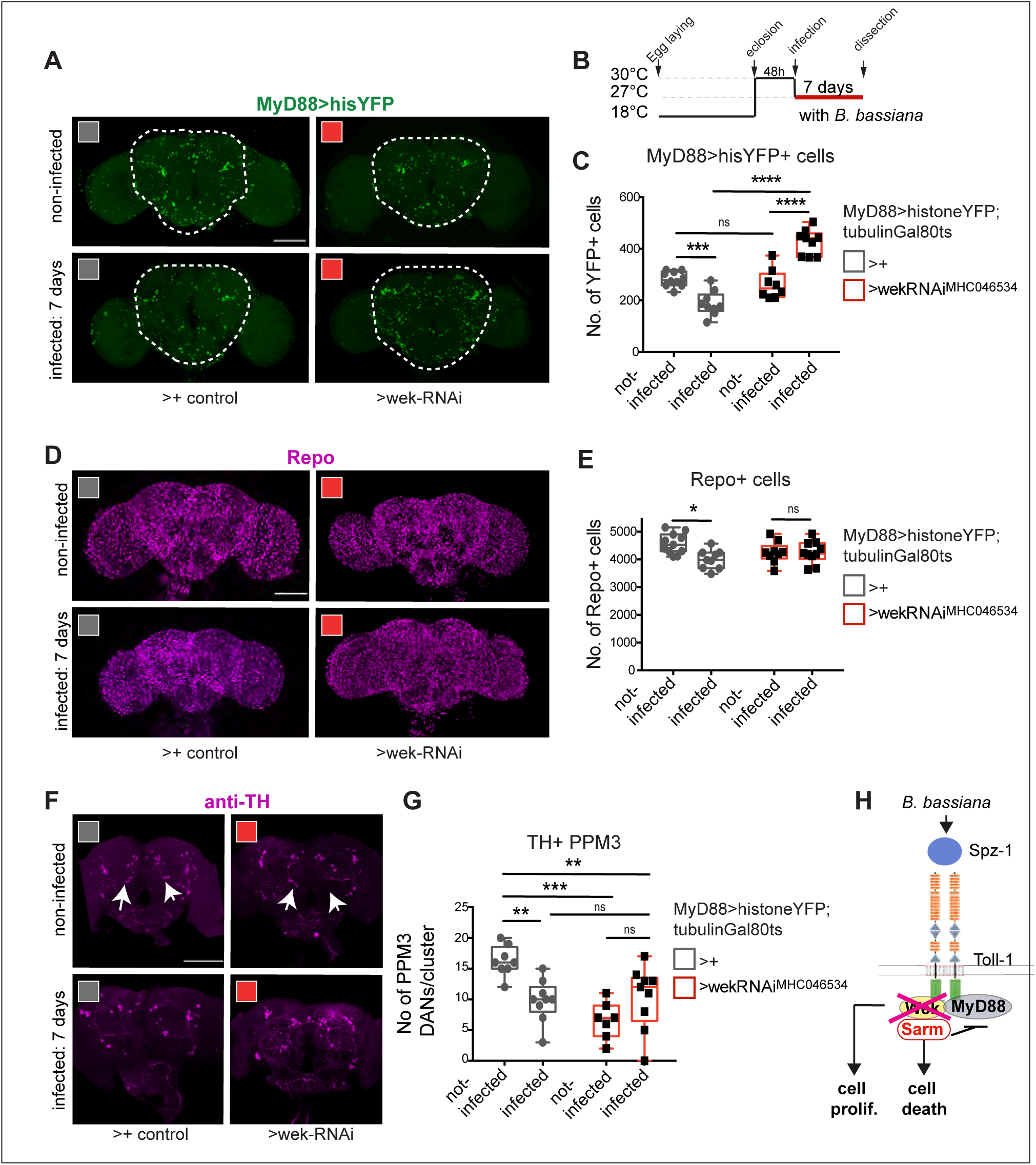
*B. bassiana* induced cell loss requires Wek, but Wek has pleiotropic functions. **(A)** Loss of *MyD88>HisYFP+* cells caused by seven-day exposure to the *B. bassiana* was rescued with adult-specific *wek-RNAi^MHC046534^*knockdown, and cell number increased significantly**. (B)** Diagram explaining the experimental regime. **(C)** Quantification of data in (A) shown as box-plots around the median. Two-way ANOVA: Infected vs not-infected p= 0.2327; Control vs wekRNAi (*tubGAL80^ts^; MyD88>HisYFP>Oregon* vs *tubGAL80^ts^; MyD88>HisYFP> wek-RNAi^MHC046534^*): p<0.0001; Interaction: p<0.0001, followed by Turkey’s multiple comparison correction test. Sample size: non-infected control brains n=14, infected control brains n=15, non-infected wek-RNAi brains n=8, infected wek-RNAi brains n=9. **(D, E)** Loss of Repo+ glial cells caused by seven-day exposure to *B. bassiana* was rescued with adult-specific *wek-RNAi^MHC046534^* knockdown, quantification in **(E).** Two-way ANOVA: Infected vs not-infected p= 0.0236; Control vs wekRNAi: p= 0.7775; Interaction: p= 0.0231, and Turkey’s multiple comparisons correction test. Sample sizes: non-infected control brains n=11, infected control brains n=9, non-infected wek-RNAi brains n=10, infected wek-RNAi brains n=10. **(F, G)** Adult-specific *wek-RNAi^JF01681^*knockdown decreased the number of TH+ PPM3 DANs, suggesting that wek may be required for their differentiation. In fact, *wek-RNAii^JF01681^* knockdown did not rescue the cell loss caused by *B. bassiana* infection, but infection did not reduce cell number further either. Two-way ANOVA: Infected vs not-infected: p= 0.3098; Control vs *wekRNAi*: p= 0.0016; Interact ion p= 0.0005, followed by Turkey’s multiple comparisons correction test: non-infected control brains n=8, infected control brains n=9, non-infected *wek-RNAi* brains n=7, infected *wek-RNAi* brains n=9. **(H)** Diagram of Toll signalling pathway, whereby *wek* links Toll-1 to Sarm, enabling Toll signalling to cause cell death and cell loss via Sarm, and this is rescued in some cells with *wek RNAi* knock-down. Wek also has functions independently of Sarm and MyD88, promoting adult neurogenesis. Dotted line indicates ROI for automatic cell counting with DeadEasy software. Asterisks on graphs: *p<0.05, **p<0.01, ***p<0.001, ****p< 0.0001. Scale bars: (A, D, F) = 100 μm.

By contrast, *wek* knock-down in non-infected adult flies caused a significant decrease in the number of DANs of the PPL1, PPL2 and PPM3 clusters (Figure 5F,G and Supplementary Figure 2). This suggested that Wek is required to maintain DAN cell survival or promote neurogenesis or differentiation (e.g. TH expression) in the adult brain. Importantly, in *wek*-RNAi knock-down flies, *B. bassiana* infection failed to induce further neuronal loss (Figure 5F,G). These data are consistent with the known pleiotropic functions of *wek,* including in the adult brain (Chen et al., 2006, Foldi et al., 2017, Li et al., 2020). They have also revealed that in the absence of Wek, some DANs do not develop normally and only a few DANs remain, but these remaining DANs are not susceptible to further damage by *B. bassiana* infection.

Next we tested *sarm.* In the absence of infection, when we downregulated *sarm* expression with RNAi in adult MyD88 cells (using line *UAS-sarm-RNAi^JF01681^*, shown to work effectively in (Foldi et al., 2017)), there was no effect compared to controls (Figure 6A-C). However, RNAi knock-down of *sarm* rescued *B. bassiana*-induced MyD88>hisYFP+ cell loss (Figure 6A-C). Similarly, *sarm* RNAi knock-down had no effect on the number of glial cells in the adult brain compared to non-infected controls, but it rescued the Repo+ cell loss caused by *B. bassiana* infection (Figure 6D,E). And finally, using anti-TH antibodies we showed that *sarm* RNAi knock-down did not affect PPM3 cell number in non-infected controls, but it rescued the loss of PPM3 DANs caused by *B. bassiana* infection (Figure 6F,G). Altogether, these data showed that *sarm* is required for *B. bassiana* induced loss of MyD88-HisYFP+, Repo+ and TH+ PPM3 cells (Figure 6H).

**Figure 6.**
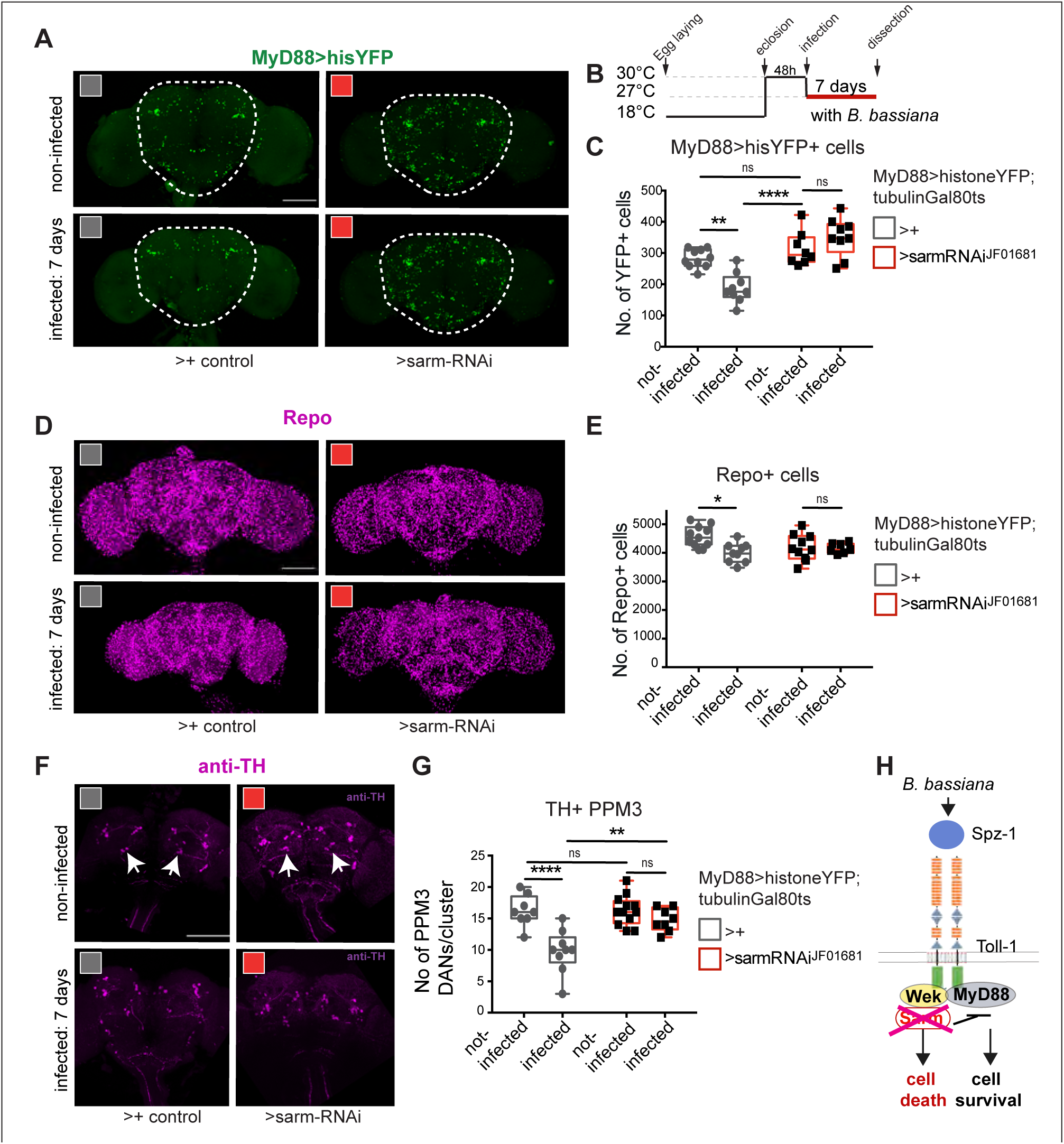
*B. bassiana* induced cell loss requires *sarm*. **(A)** Seven-day exposure to the *B. bassiana* caused loss of *MyD88>HisYFP+* cells, and this was rescued with adult-specific *sarm-RNAi^JF01681^*knockdown, quantification in (C). **(B)** Diagram explaining the experimental regime. **(C)** Quantification of data in (A), Two-way ANOVA: Infected vs not-infected p= 0.0077; Control vs sarmRNAi (*tubGAL80^ts^; MyD88>HisYFP>Oregon* vs *tubGAL80^ts^; MyD88>HisYFP> sarm-RNAi^JF01681^*): p<0.0001 Interaction: p<0.0001, followed by Turkey’s multiple comparison correction test. Sample sizes: non-infected control brains n=14, infected control brains n=15, non-infected sarm-RNAi brains n=8, infected sarm-RNAi brains n=9. **(D, E)** Seven-day exposure to the *B. bassiana* caused loss of Repo+ glial cells **(D),** and this effect was rescued with adult-specific *sarm-RNAi^JF01681^* knockdown, quantification in **(E).** Two-way ANOVA: Infected vs not-infected p= 0.0205; Control vs *sarmRNAi*: p= 0.3649; Interaction: p= 0.0295, followed by Turkey’s multiple comparisons correction test. Sample sizes: non-infected control brains n=11, infected control brains n=9, non-infected *sarm-RNAi* brains n=10, infected *sarm-RNAi* brains n=7. **(F, G)** Seven-day exposure to *B. bassiana* caused loss of TH+ PPM3 DANs, which was rescued with adult-specific *sarm-RNAi^JF01681^* knockdown, quantification in **(G)**. Two-way ANOVA: Infected vs not-infected p< 0.0001; Control vs *sarmRNAi*: p= 0.0071; Interaction: p= 0.0071, followed by Turkey’s multiple comparisons correction test. Sample sizes: non-infected control brains n=8, infected control brains n=9, non-infected *sarm-RNAi* brains n=11, infected *sarm-RNAi* brains n=8. **(H)** Diagram of Toll signalling pathway, whereby Sarm leads to cell death and cell loss. B. bassiana infection caused cell-loss is rescued with *sarm* RNAi knock-down. Dotted line indicates ROI for automatic cell counting with DeadEasy software. Data shown as box-plots around the median. Asterisks on graphs indicate multiple comparisons corrections: *p<0.05, **p<0.01, ****p< 0.0001. Scale bars: (A, D, F) = 100 μm.

Altogether, these data show that signalling via Wek and Sarm is required for *B. bassiana*-induced cell loss in the brain. This suggests that the upregulation of *wek* and *sarm* expression in the brain upon *B. bassiana* infection could lead to neurodegeneration in the host (Figure 2F and Figure 6H).

### Increased Toll-1, Wek and Sarm levels could induce cell loss

Finally, we asked whether over-expression of *activated Toll-1, wek* or *sarm* could be sufficient to induce cell loss in the absence of infection. Wek has pleiotropic functions downstream of Tolls: it binds Tolls and MyD88 to enable canonical signalling via NFkB/Dorsal/Dif downstream, which in the CNS promotes quiescence and cell survival; it can also bind Sarm, to promote cell death; or function independently of both, to promote neurogenesis in the adult brain from quiescent MyD88+ progenitor cells (Chen et al., 2006, Foldi et al., 2017, Li et al., 2020). Thus, the pleiotropic functions of *wek* could lead to compound phenotypes. To simplify this, we tested the effect of activated *Toll-1^10b^*, *wek* or *sarm* over-expression on PAM neurons, which are differentiated neurons. We used a DAN specific GAL4 driver (*THGAL4; R58E02GAL4*), visualised PAMs with histone-YFP and counted them automatically. Toll-1 is normally expressed only in a fraction of PAM neurons (Supplementary Figure 1). Over-expressed activated *Toll-10^1b^* in DANs caused a mild decrease in PAM cell number, which however was not statistically significantly different from the control (Figure 7A,B). This suggested that not many PAMs may express *wek, sarm* or both in the un-infected brain. Importantly, over-expression of either *wek* (Figure 7A,B) or *sarm* (Figure 7C,D) was sufficient to cause a significant reduction in the number of PAM neurons in the absence of infection. This shows that increased Wek and Sarm levels could induce PAM cell loss.

**Figure 7.**
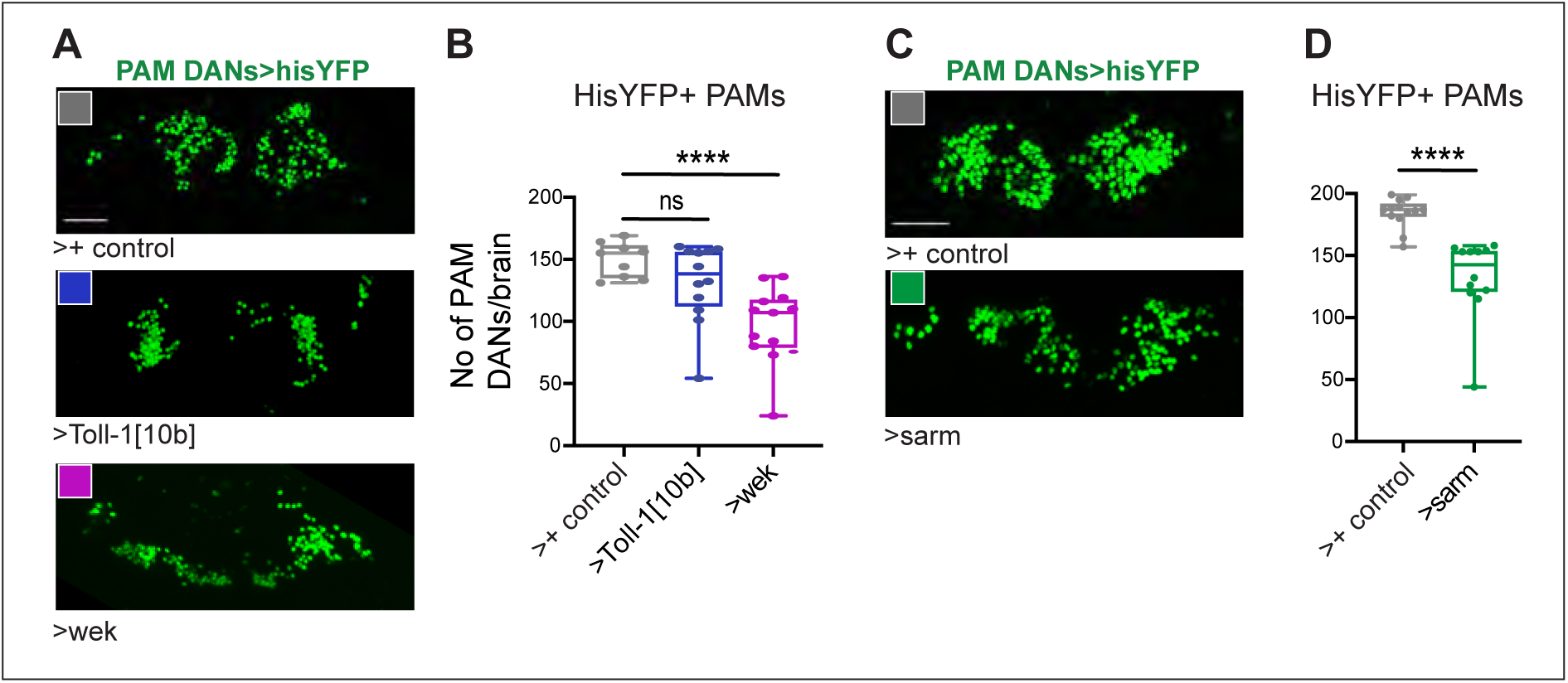
Increased *Toll-1, wek* and *sarm* levels can induce cell loss. **(A,B)** In the absence of infection, over-expression of activated *Toll-1^10b^*in DANs (with *THGAL4; R58E02GAL4*) caused a rather mild and not significant decrease in PAMs. By contrast, over-expression of *wek* was sufficient to induce cell loss (7-day old flies). Sample sizes: control n=9 brains, *UAS-Toll-1^10b^* n=12*; UAS-wek-HA* n=13. One-Way ANOVA p=0.0003, multiple comparisons corrections Dunnett test to a fixed control. **(C,D)** Over-expression of *sarm* was sufficient to induce PAM cell loss in the absence of infection (2-day old flies). Flies were kept constantly at 25°C. Student t test. Asterisks on graphs indicate multiple comparisons corrections: ****p< 0.0001. Scale bars: (A,C) 30μm.

Altogether, these data support the notion that Toll-1 signalling via Wek and Sarm can induce cell loss in the brain. They indicate that the upregulation of *wek* and *sarm* caused by *B. bassiana* infection leads to cell loss in the host brain.

## DISCUSSION

We show here that the fungus *B. bassiana* induces cell loss in the host *Drosophila* brain via the innate immunity Toll-1 receptor. In innate immunity, Toll-signalling via MyD88 and NFkB/Dif results in the upregulation of anti-microbial peptide gene expression to eliminate the fungus (Imler and Hoffmann, 2001, Lemaitre et al., 1996, Leulier and Lemaitre, 2008). However, we show that in the brain, *B. bassiana* can also activate an alternative Toll-1 signalling pathway, involving Wek and Sarm, that causes neurodegeneration instead. Sarm is the highly evolutionarily conserved and only inhibitor of Toll/TLR-dependent innate immunity, from flies to humans (Anthoney et al., 2018, Belinda et al., 2008, Carty and Bowie, 2019, Peng et al., 2010, Yuan et al., 2010). Through its ability to up-regulate Sarm, *B. bassiana* drives immune evasion, causing neurodegeneration in the host brain.

*B. bassiana* spores infiltrated the brain, damaged the BBB and upregulated the expression of antifungal peptides *drs* and *mtk* in infected brains. *B. bassiana* might enter the brain by first adhering to and degrading the cuticle (Boucias et al., 1988), in the optic lobes, head capsule and proboscis. Once attached, *B. bassiana* spores germinate and produce hyphae that degrade the cuticle by secreting various chitinases and proteases (Fan et al., 2007, Fang et al., 2009, Zhang et al., 2008). Spores in the brain and a compromised blood-brain barrier were also found following *E. muscae* infection in *Drosophila* (Elya et al., 2023, Elya et al., 2018). *B. bassiana* could also trigger a host response facilitating macrophage entry across the BBB and into the brain (see (Winkler et al., 2021)). In many insects, the likelihood of infection is highest at less sclerotized cuticle regions like the mouth (Ortiz-Urquiza and Keyhani, 2016). As the proboscis traverses the brain, dissolving the proboscis cuticle would provide *B. bassiana* a direct route to the brain. Similarly, fungi also invade the human brain (Cannon and Gruenheid, 2022, Chen et al., 2022, Pisa et al., 2020, Pisa et al., 2015a, Pisa et al., 2015b). The most common infection is by *Cryptococcus*, which causes meningoencephalitis, and *Cryptococcus* enters the human brain through the nose, by inhalation, followed by degradation of the BBB (Chen et al., 2022).

We showed that exposure to *B. bassiana* decreased fly longevity and impaired locomotion, which correlated with widespread cell loss in the brain. *B. bassiana* induced-cell loss was verified with multiple cell markers: loss of His-YFP labelled Sarm+ and MyD88+ cells, loss of Repo+ glial cells and loss of TH+ DANs. Furthermore, *B. bassiana* infection reduced the expression of TH, which is responsible for dopamine synthesis, and dopamine is required for locomotion (Riemensperger et al., 2013). Our findings are consistent with reports that fungal volatiles induce neurodegeneration in the *Drosophila* brain (Inamdar et al., 2013, Inamdar et al., 2010). Evidence of neurodegeneration were decreased longevity, impaired climbing, decreased dopamine levels and dopaminergic neuron loss (Inamdar et al., 2013, Inamdar et al., 2010). These same phenotypes are also measures of neurodegeneration in other contexts, such as *Drosophila* models of Parkinson’s disease (Riemensperger et al., 2013, Cassar et al., 2015, Feany and Bender, 2000). Similarly, our work demonstrates that exposure to *B. bassiana* reduced longevity, impaired locomotion, caused cell loss – including of DANs – and reduced TH levels. Altogether, exposure to *B. bassiana* induced a signature characteristic of neurodegeneration. Notably, *B. bassiana* is also a plant endophyte, providing protection to plants against insects (Behie et al., 2012, Behie et al., 2015, Branine et al., 2019). By reducing the climbing ability of flies, *B. bassiana* may facilitate greater spore adherence and germination on the fly cuticle, ultimately leading to the insect’s demise.

*B. bassiana* induced neurodegeneration involved signalling via the Toll-1 immune-evasion pathway, involving Wek and Sarm. Toll-1 cannot directly bind Sarm, and instead it binds Wek at the plasma membrane, and Wek binds Sarm (Chen et al., 2006, Foldi et al., 2017). In this way, Wek enables Toll signalling via Sarm (Foldi et al., 2017). *B. bassiana* infection caused an increase in *wek* and *sarm* expression within the brain, opening this alternative signalling option downstream of Toll-1. Similarly, *E. muscae* infection also induced the upregulation of *wek* (Elya et al., 2018). Importantly, knocking down either *Toll-1, sarm* or *wek* rescued the cell loss caused by *B. bassiana* infection, and over-expression of *wek* or *sarm* was sufficient to induce cell loss in the absence of infection. Together, these data demonstrate that *B. bassiana*-induced cell loss requires Toll signalling via Wek and Sarm. Importantly, we showed that although Toll-1 expression varies among different DAN clusters, Toll-1 knock-down rescued *B. bassiana*-induced loss of cells expressing Toll-1. The variable effects in distinct DANs also suggest that different DAN clusters may express different combinations of Tolls, downstream adaptors and effectors driving distinct outcomes. In fact, different Tolls can have distinct effects on survival or death (Foldi et al., 2017). For example, at least Toll-1, Toll-2,-6 and-7 can induce cell survival via MyD88 (Li et al., 2020, McIlroy et al., 2013, Zhang et al., 2024, Zhu et al., 2008); Toll-2 induces neurogenesis via *wek* (Li et al., 2020); and Toll-1,-6 and-7 can induce cell death when co-expressing *MyD88, wek* and *sarm* (Foldi et al., 2017). It has been proposed that Toll-1 and-7 maintain DAN cell survival via regulating autophagy, through MyD88 (Zhang et al., 2024). This suggests that PAMs normally express MyD88. By contrast, we found that *Toll-1^10b^*could not induce severe PAM cell loss, which could be explained if not all PAMs express both *wek* and *sarm*. Absence of Wek in PAMs would prevent Toll-1 from driving apoptosis, even in the presence of Sarm, as Tolls cannot bind Sarm without Wek (Foldi et al., 2017). Instead, over-expression of *wek* or sarm was sufficient to induce PAM cell loss. On the other hand, some DANs (PPL1, PPL2 and PPM3) were lost with *wek* knock-down, meaning that *wek* is required for their cell survival, neurogenesis or differentiation. This is consistent with the fact that *Toll-2* promotes adult neurogenesis via Wek (Li et al., 2020). It is currently unknown if *wek* can directly regulate neuronal differentiation, but it is not unlikely, as Wek encodes a zinc-finger transcription factor (Chen et al., 2006). Either way, the data further support the notion that Wek has pleiotropic functions that depend on cellular context (Foldi et al., 2017, Chen et al., 2006).

The data also suggest that signalling by Toll-1/Wek/Sarm may not be enough to induce widespread neurodegeneration after infection. One possibility is that other Tolls might also induce cell death when Wek and Sarm levels rise. In fact, Spz-1 can bind at least Toll-7 in flies (Chowdhury et al., 2019), and Toll-5 in mosquito (Saucereau et al., 2022).

Alternatively, anti-microbial peptides induced by Toll-1 via MyD88/NFkB/Dif signalling could also drive neurodegeneration non-autonomously. In fact, anti-microbial peptides regulated by NFκB Rel downstream of the Imd pathway can induce neurodegeneration (Kounatidis et al., 2017), and over-expression of both *drs* and *mtk* can induce DAN cell loss (Shukla et al., 2019). In *C. elegans*, TIR-1/Sarm promotes the expression of anti-microbial peptides in the epidermis, which bind a receptor in neurons to cause dendrite degeneration (E et al., 2018). Thus, *B. bassiana*’s ability to induce neurodegeneration in the host may involve a wider molecular machinery than explored here.

Our data suggest that the activation of Sarm signalling by *B. bassiana* is one of a range of adaptive interactions with its hosts. For example, in mosquitos, *B. bassiana* can reduce the host immune response by exporting a microRNA that downregulates the levels of Spz4, a ligand of Toll (Cui et al., 2019). In flies, *B. bassiana* can bypass the host pattern recognition receptor GNBP3 by releasing virulence factors that promote host immune evasion and fungal growth within the host (Eley et al., 2007, Molnár et al., 2010, Pedrini, 2022). In return, flies can detect the virulence factors too, to activate innate immunity bypassing detection of the fungal cell wall through GNBP3. In fact, the virulence factor PR1 gets proteolytically cleaved by Persephone, and activates the Toll-signalling cascade independently of GNBP3 (Gottar et al., 2006). Similarly, in mammals, Toll-Like Receptor-4 (TLR-4) activates the immune response against *C. albicans,* while *C. albicans* activates TLR-2, which leads to production of anti-inflammatory chemokines that help *C. albicans* to evade the host immune system (Netea et al., 2004, Netea et al., 2002, Netea et al., 2007). In this context, our data show that *B. bassiana* can induce the Sarm pathway downstream of Toll-1 harming the host. What controls the up-regulation of *wek* and *sarm* after infection, enabling degenerative signalling, is unknown, but it could be Toll signalling itself, activated by *B. bassiana*. In fact, signalling downstream of Toll-6 and-7 upregulates the expression of various Toll downstream effectors, including *NFκB* homologues *dorsal* and *dif,* and their inhibitor *cactus* (McIlroy et al., 2013). Importantly, the functions of Sarm in inhibiting canonical Toll/TLR signalling and in inducing neuronal apoptosis, axon destruction and neurodegeneration are all highly evolutionarily conserved (Anthoney et al., 2018, Belinda et al., 2008, Carty and Bowie, 2019, Sarkar et al., 2023). In flies and mammals, Sarm induces apoptosis via JNK signalling and in *C. elegans*, via MAPK/p38 signalling (Carty et al., 2006, Essuman et al., 2017, Izadifar et al., 2021, Mukherjee et al., 2015, Osterloh et al., 2012, Peng et al., 2010, Veriepe et al., 2015, Yuan et al., 2010, Foldi et al., 2017). Moreover, Sarm has a TIR domain that has catalytic NADase activity, which drives neurite destruction as well as cell death (Essuman et al., 2017, Osterloh et al., 2012, Summers et al., 2016). Most intriguingly, similarly to Sarm, some prokaryotic proteins also bear a TIR domain with NADase catalytic function, and they function in immune evasion (Sarkar et al., 2023).

To conclude, our data show a novel tactic in the evolutionary arms race between the host *Drosophila* and the fungus *B. bassiana* taking place in the brain. *B. bassiana* is detected in the *Drosophila* brain, that activates Toll-1 dependent innate immunity signalling against the fungus to protect the brain. In turn, *B. bassiana* benefits from the concomitant activation of the Toll-1 immune-evasion Sarm pathway driving neurodegeneration in the host brain. Importantly, human neurodegenerative and psychiatric diseases have been linked to fungal infections and neuro-inflammation (Chen et al., 2022, Alonso et al., 2014, Pisa et al., 2020, Cannon and Gruenheid, 2022). There is also evidence linking altered Sarm function to neurodegenerative diseases (Bloom et al., 2022, Miao et al., 2024, Veriepe et al., 2015, Murata et al., 2023). For example, constitutively active Sarm1 variants are enriched in ALS patients (Bloom et al., 2022). It will be important to find out whether a similar activation of Sarm downstream of TLRs in response to fungal infections is responsible for inducing psychiatric and neurodegenerative diseases in humans.

## MATERIALS AND METHODS

### Drosophila genetics

Please see Table 2.1 for the list of the stocks used. Conditional over-expression and knock-down in adult flies was carried out using ubiquitously expressed *tub-GAL80^ts^*. *GAL80^ts^* is a temperature sensitive GAL4 repressor, that prevents GAL4 expression at 18°C and enables it at 30°C. The temperature regimes we used are indicated in the figures and were set to enable GAL4 adult-onset and *B. bassiana* growth together.

**Table 1.**
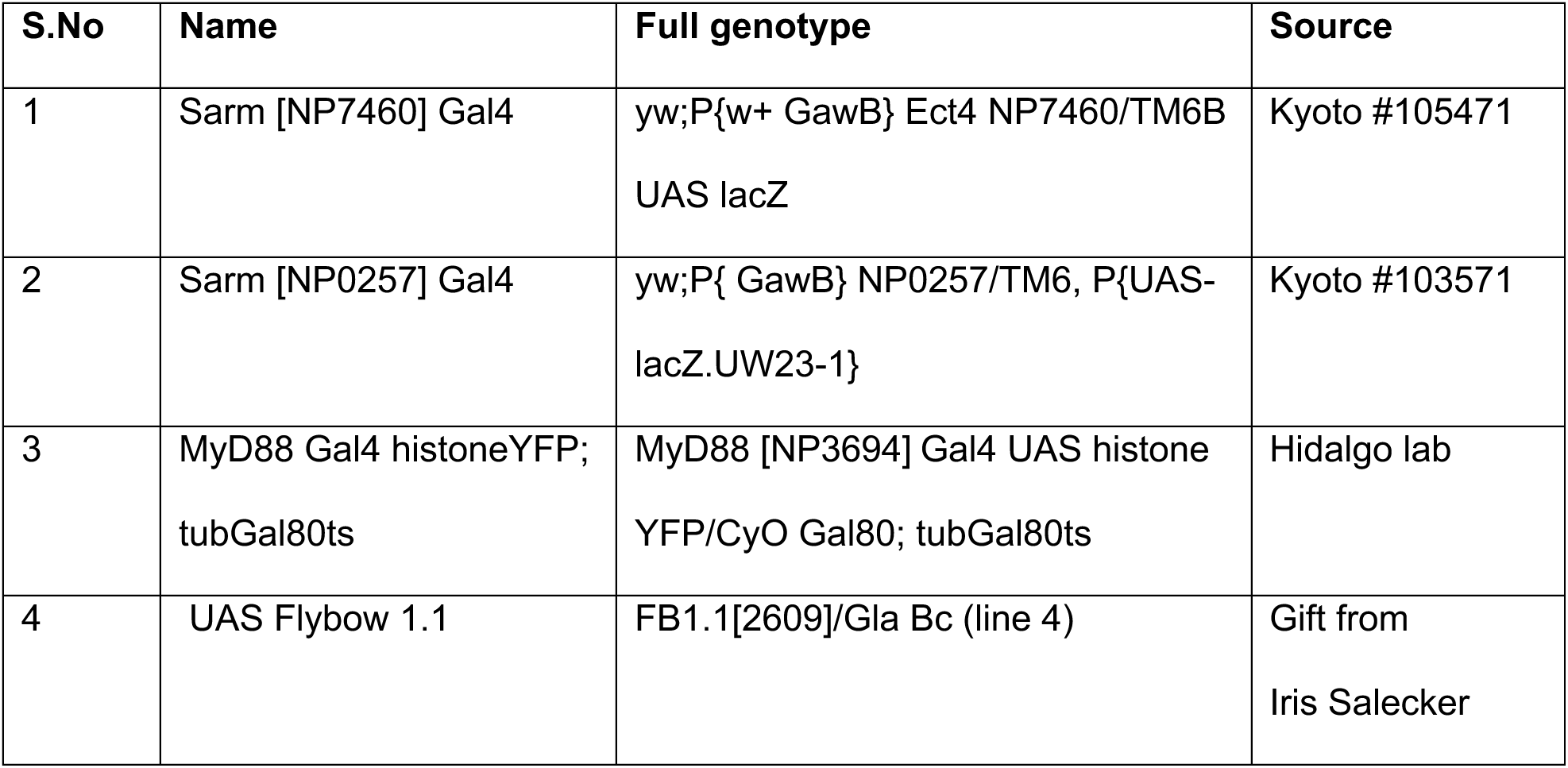
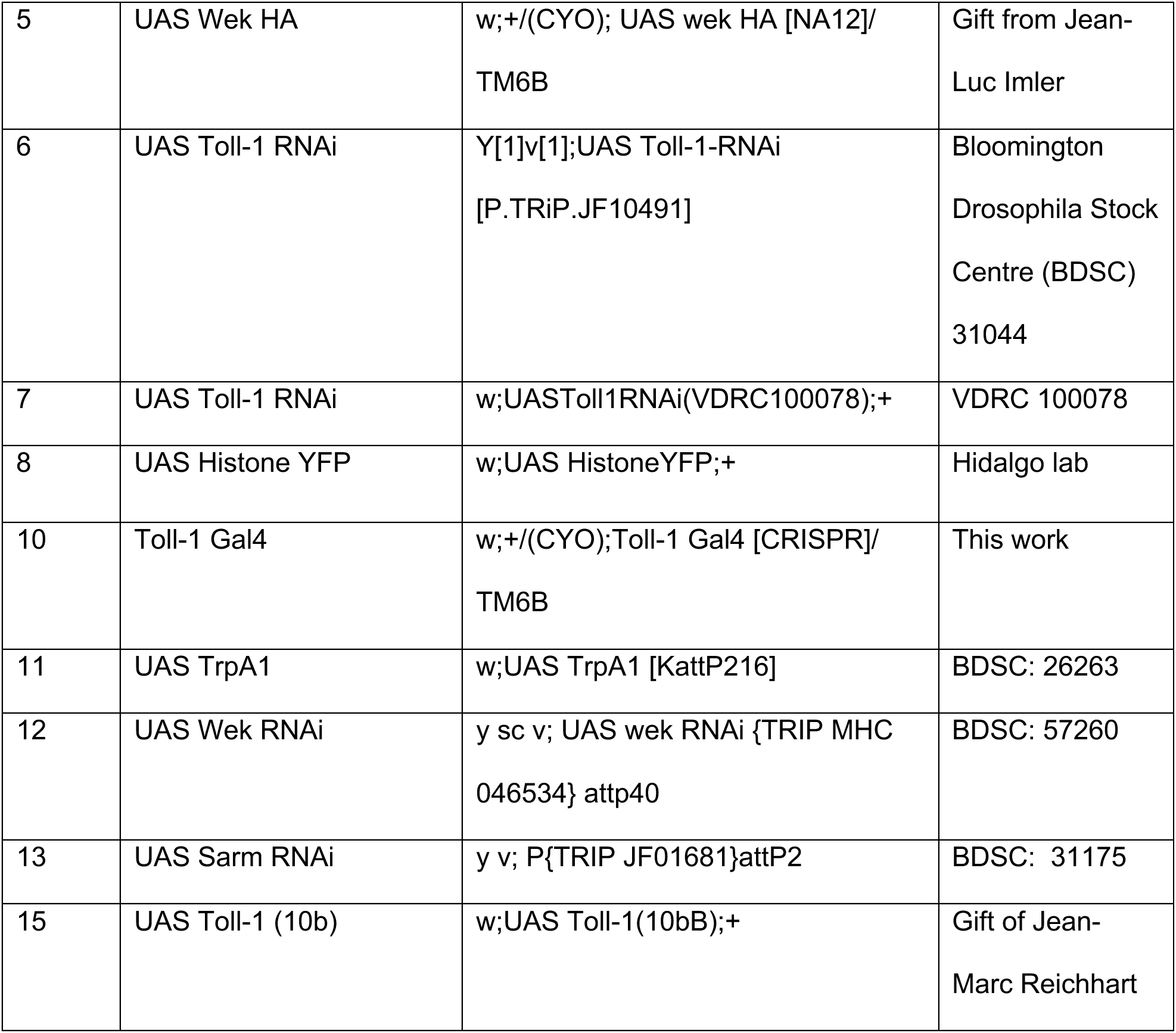
Stock list.

**Table 2.**
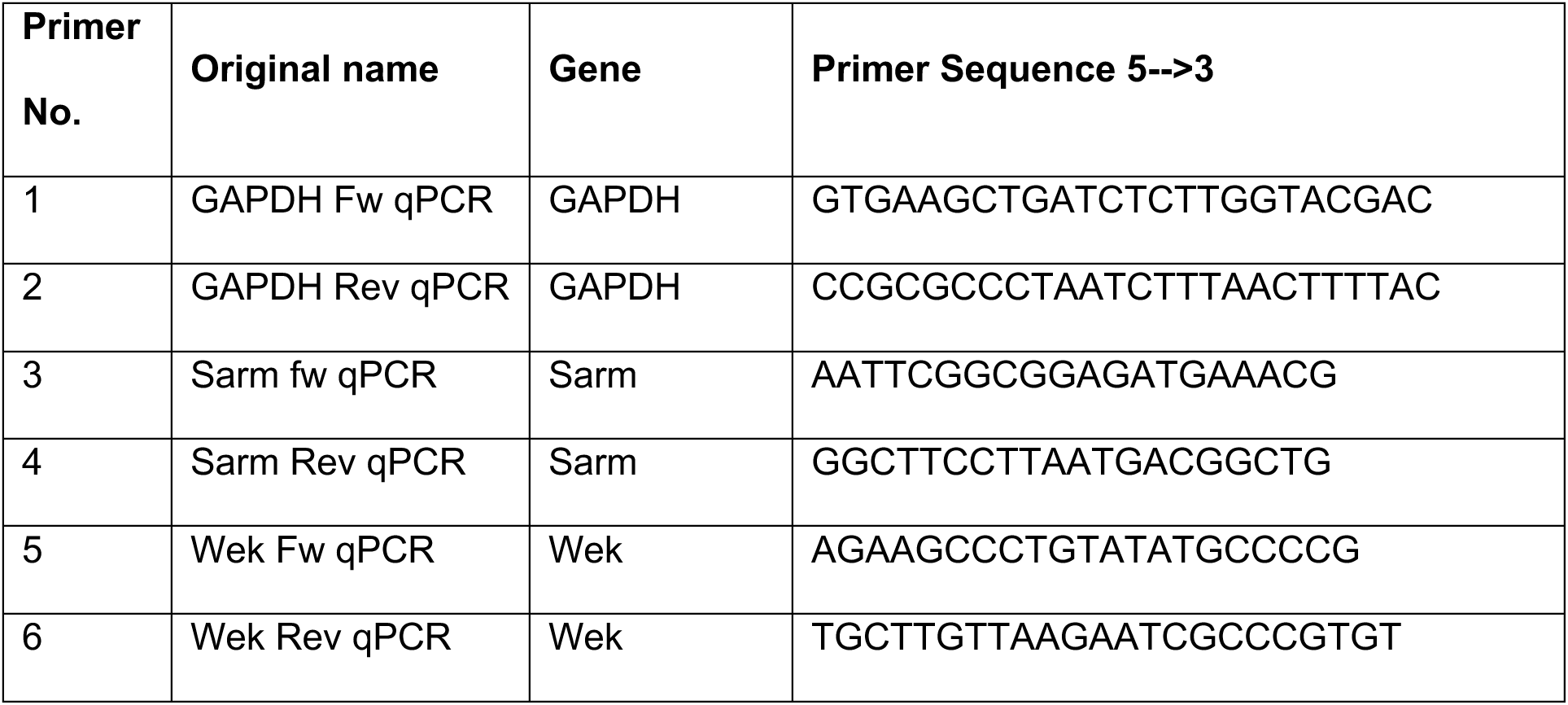
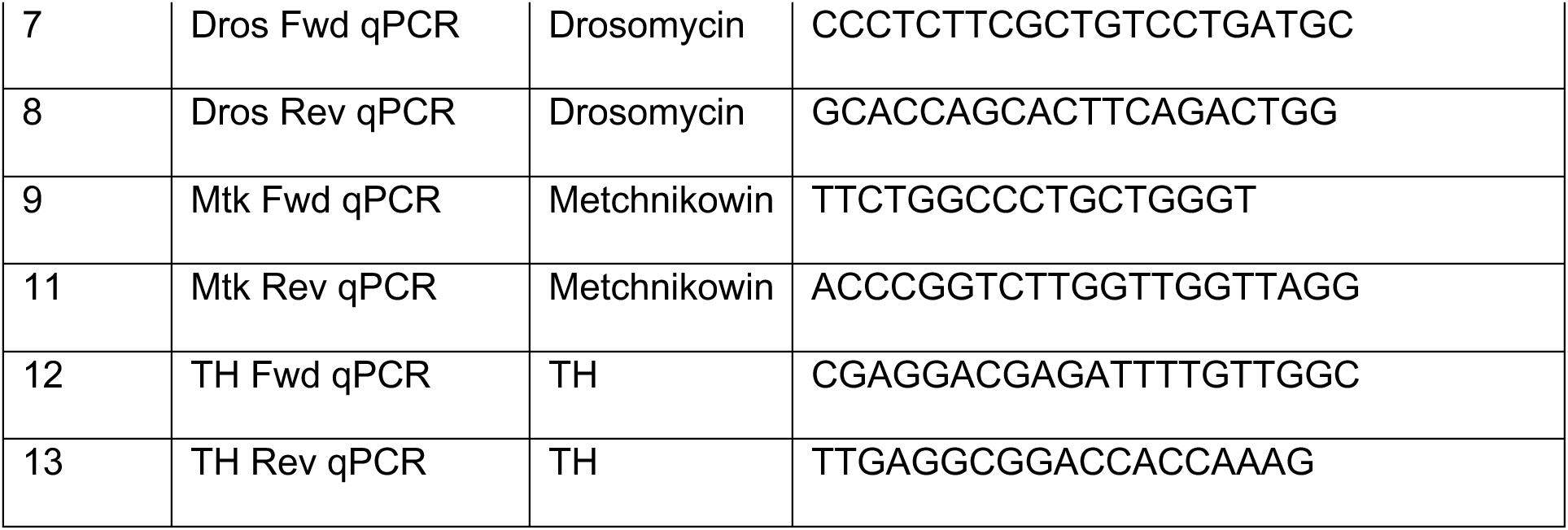
List of primers.

### *B. bassiana* culture, spore collection and infection chambers

*Beauveria bassiana* (80.2 strain) was a gift from Jean-Marc Reichhart and Jean-Luc Imler, (IBMC/University of Strasbourg); GFP transgenic *B. bassiana* (EABb 04/01-Tip GFP5 strain) was a gift from Prof. Enrique Quesada-Moraga (University of Cordoba).

#### Spores

To isolate *B. bassiana* spores, 10 ml of distilled water was poured on the *B. bassiana* culture in the petri plate and culture was scraped with a spreader. The solution was centrifuged at 4,000 rpm for 15mins, the supernatant was removed, the pellet was dissolved in 2-5ml of water and spun again for 1-2mins. The pellet was dissolved in 1ml distilled water and labelled as the principal solution. The concentration of the principal solution was calculated by counting spores using a haemocytometer. 10µl of principal solution was diluted in 90µl of water and marked as dilution X10 and spores were counted again by using the haemocytometer. This step was repeated until a spore concentration of 3.7 x 10^9^ spores/ml was achieved.

#### Infection chamber

To enable natural infection whilst avoiding damage to the body that would induce an injury-response, a natural infection chamber was devised. An infection chamber consists of a glass bottle with a carpet of *B. bassiana* growing on sabouraud dextrose agar (SDA) medium at the bottom, and a cut 50ml falcon tube bearing standard fly-food stuck with double-sided sticky tape to the bottle side wall. To prepare SDA media, 16.25g of 4% SDA was added to 250 ml of distilled water and autoclaved. Next, 10ml SDA medium was transferred into autoclaved glass bottles and once it had set, 1ml of 3.7 x 10^9^ spores/ml *B. bassiana* solution was poured into each bottle onto the medium. After plugging the bottles, they were transferred to 25°C incubator for 7-10 days. Next, the falcon tube containing fly-food was inserted into the bottle, and then the flies. In this way, flies can freely access fly-food devoid of *B. bassiana*, for feeding and hydration, whilst being naturally exposed to spores.

### Staining of *B. bassiana* spores

Adult Drosophila brains were dissected and fixed in 4% paraformaldehyde (PFA) in Phosphate buffer saline (PBS) at room temperature for 20 minutes. Following washes in 0.5% Triton in PBS, brains were incubated in FM4-64 dye at 1:200 dilution, or calcofluor white (CLW) stain (50mg/ml) 1:1000 dilution for 1h at room temperature. The brains were washed and mounted in DAPI containing Vectashield, prior to imaging.

### Drosophila behaviour tests

**Longevity** was measured as described in (Piper and Partridge, 2016), using F1 wild-type Oregon/CantonS flies raised on standard cornmeal food. They were placed in an infection chamber, the fly food was replaced every four days in both control and experimental setups, and the number of dead flies and censor events were manually recorded after every transfer. Both infected flies and non-infected controls were kept at 25°C. Dead flies were scored as 1 and flies that got stuck to the food or escaped were scored as 0. Data were analysed using GraphPad prism and Log rank (Mantel-Cox test) to analyse the data and generate a Kaplan-Meier survival curve. The experiment was conducted with n=104 flies for each condition, from two biological replicates of n=58 and n=46 flies each.

### Climbing assay

Startled induced negative geotaxis assay was carried out as described in (Barone and Bohmann, 2013). The test was carried out in a humidity and temperature controlled lab (25°C). The infected flies were flipped 2-3 times to remove excess spores attached to their bodies before being transferred to climbing assay vials. The vials with flies inside were tapped, and the flies were filmed for 10 seconds, followed by a 30-second rest period, and the number of flies climbing above the 2cm mark were counted. This process was repeated 15 times for each cohort. Three cohorts of 7-10 flies each were used per genotype, and the entire process was repeated a second time, using new flies.

### Proboscis extension response (PER) assay

The PER assay was adapted from (Smith and Burden, 2014). Flies were starved in a 25°C incubator for 24 hours in vials containing agar before performing the PER assay. After 24hrs, the flies were transferred to a 20µl pipette tip using a fly aspirator. Using a sharp razor blade, the tip was cut, and the fly was then carefully placed in position with the help of a wick made of Kimtech wipe. The 20µl tip with the fly head emerging was fixed on a flat surface and a camera was positioned around it. To verify that the immobilized flies in the 20µl tip were fit for the PER assay, they were first given water and 100 mM sucrose solution using tissue paper wicks and the response was recorded for 60 seconds. The flies had been hydrated but starved before the experiment, so any flies that responded to water or did not respond to the 100 mM sucrose solution were discarded. The TrpA1 experiment was conducted inside an environmental chamber at 30°C. After the flies had been inside the chamber for 2-3 minutes, they were recorded for 60 seconds. The experiment was conducted using 10 flies per genotype, and the process was repeated three times using new biological replicates of the crosses.

### Feeding assay

The feeding assay was adapted from (Cheriyamkunnel et al., 2021). To prepare a fly cage, an empty food vial was used, and was cut at 4mm from its base using a heated knife. The base was then sterilized with 70% ethanol. Solutions mixed with 4% brilliant blue dye were transferred to the base and placed in an egg-laying chamber. For the feeding assay, four solutions were prepared, with each egg laying chamber containing only one solution: Negative control 1: distilled H2O; Negative control 2: distilled H2O (950ul) + 50ul 4% brilliant blue dye (95%/5% v/v); Positive control: 100mM Sucrose solution (950ul) + 50ul 4% brilliant blue dye (95%/5% v/v); Test: 7.0025 x 10^7^ spores/ml (950ul) + 50ul 4% brilliant blue dye (95%/5% v/v). Equal number of wildtype Oregon male and female flies were used (n=10 males and 10 females) and 4 biological replicates were performed, flies were anesthetized on ice and transferred to the 4-feeding assay setups containing the above solutions. The set-ups were then placed at 25°C incubator for 24hrs. Next, the flies were collected by anesthetising the flies with CO2 and transferred to the 1ml Eppendorf tube. The Eppendorf containing flies were dipped in liquid nitrogen for 5 secs, placed in the 50ml Falcon tube, and dropped from a height to separate the flies’ heads from their abdomens. The abdomen of the flies was collected using forceps and transferred to an Eppendorf containing 50ul of distilled water. Using a pestle abdomen was homogenised in the Eppendorf and 950ul of dH2O was added to it. The Eppendorf was then vortexed and centrifuged for 5 mins at 10,000rpm. The supernatant was transferred into the new Eppendorf and kept on ice until analysis. The homogenised solution was transferred to 1ml cuvette, and the absorbance was recorded using spectrophotometer, at 630nm and 750nm. The corrected absorbance was calculated by subtracting the mean of 3 readings at 750nm (background) from the mean of 3 readings at 630nm (peak of blue dye).

### Injecting flies with Dextran red fluorescent dye

This protocol was adapted from (Bainton et al., 2005). A glass pulled capillary needle was inserted into a glass pasteur pipette and sealed with parafilm at the point of insertion, and needle was loaded with 0.1µl of 25 mg/µl fluorescent Dextran red dye. The needle was carefully inserted into the thorax beneath the wing of anaesthetised flies. The flies were then transferred to a vial containing fly food and placed in a 25°C incubator for 24hrs to recover. After the 24hrs recovery period, the flies were fixed onto a glass slide using glue, and their retinas were imaged using Leica SP8 confocal fluorescent microscope, n=10 flies.

### Quantitative Reverse Transcription (qRT)-PCR

qRT-PCR was carried out from 20 adult brains per sample, dissected from wild type Oregon non-infected and infected flies. The dissected adult brains were cleaned removing all fat body and then immediately transferred into the Eppendorf containing TRI reagent (Ambion #AM9738). RNA extraction was carried out according to the TRI reagent protocol using isopropanol. Extracted RNA were treated with DNase to get rid of genomic DNA contamination using DNA-free kit (Ambion #AM1906). 200ng of RNA was reverse transcribed into cDNA by using random primer by following the manuscript of GoScript^TM^ Reverse Transcription System (Promega #A50001) (Table 2.1). A PCR was performed to check the contamination in the RNA and Reverse Transcribed (RT) cDNA (Table 2.2). This is followed by a qRT-PCR. All qRT-PCR experiments were done in triplicate (i.e., 3 well condition). A DNA binding dye SYBR was used to observe the dynamics of the PCR (SensiFast^TM^ SYBR®) and used ABI Prism 7000SDS machine for the qPCR (Table 2.3). Following qRT-PCR, quantification were performed using the CT value generated by the qPCR machine, as described in (Rao et al., 2013). For the internal control GAPDH (housekeeping\reference gene) was used. The expression of the target gene (ΔCT) was normalised in infected and non-infected flies by subtracting the CT value of the GAPDH (reference gene) from the CT value of the target gene. ΔΔCT was calculated by subtracting the ΔCT (Target gene) of the infected flies from the ΔCT (Target gene) of the non-infected flies. The relative gene expression of the target gene was then calculated in infected and non-infected flies and expressed as fold change (2^-ΔΔCT^). The statistical analysis was conducted on the ΔCT values. 3 biological replicates were used each biological replicate consist of 20 flies. ΔCT = CT (a target gene) – CT (a reference gene\GAPDH); ΔΔCT = ΔCT (non-infected sample) – ΔCT (infected sample); Fold change = 2^-ΔΔCT^

### Generation of Toll-Gal4 flies

Toll-1-GAL4 transgenic flies were generated using CRISPR/Cas9 enahnced homologous recombination. The 5’ homology arm was PCRed using primers 5’Fwd: ATGCGACCGGTAAAATCTCGTATTATGCAGCACTCGA and 5’Rev: GGAACTGAGCGGCCGCTGCAAATGGAGAAATTGAAAGGAAT; the 3’ homology arm was PCRed using primers 3’Fwd: GATGGCGCGCCGTGAACCCATTTGGACAACA and 3’ Rev: CGTACTAGTGCAGTTCAGCTCTCAGCCGT. The donor vector was pT-GEM-T2A-Gal4, and homology arms were cloned into AgeI and NotI (5’ homology arm) and AscI and SpeI (3’ homology arm) enzyme sites (as in (Gratz et al., 2015)). The guide RNA targeted the start of the coding sequence: sense oligo: GTCGCCCATTTGGACAACATGAGTCGA; antisense oligo: AAACTCGACTCATGTTGTCCAAATGGG. The gRNA was cloned into the BbsI site of vector pU6.3, as decribed in (Port et al., 2014). Thansgenesis was carried out by the BestGene. 3xP3-DsRed was subsequently removed with CRE-recombinase using conventional genetics.

### Immunostainings

Adult fly brains were dissected and fixed following standard methods, as in (Li et al., 2020). Antibodies used, their source and their working dilutions are given in Table 3.

**Table 3.** List of antibodies.

| Antibody | Donor | Working dilution |
| --- | --- | --- |
| **Primary antibodies** |  |  |
| Anti-GFP | Rabbit | 1:250 |
| Anti-REPO | Mouse | 1:20 |
| Anti-ELAV | Rat | 1:250 |
| Anti-TH | Rabbit | 1:250 |
| **Secondary antibodies** |  |  |
| Anti-rabbit-488 | Donkey | 1:500 |
| Anti-mouse-546 | Goat | 1:500 |
| Anti-mouse-488 | Goat | 1:500 |
| Anti-rat-647 | Goat | 1:500 |

### Microscopy and imaging

*B. bassiana* spores in the adult brain stained with FM4-64 dye, Calcofluor white stain, and transgenic *B. bassiana*-GFP spores were imaged with Leica SP8 confocal microscope with a 20X oil immersion lens, at a resolution of 1024×1024 pixels, zoom 1 and scanning speed of 400 Hz, and 0.96 µm Z step. High magnification images of *B. bassiana* spores in adult brain were obtained using a 63X oil immersion lens at 1024×1024, zoom 2.0, speed of 400Hz, line average of 4 and Z step 0.96 µm.

Diffusion of Dextran Red dye in the retina of adult flies were imaged using Leica SP8 confocal microscope with a 10X lens, at a resolution of 1024×1024 pixels. The zoom was set to 0.8 and scanning speed was 400Hz.

Toll-1>FB1.1, Sarm>FB1.1 proboscis were imaged using Zeiss LSM Airyscan confocal microscope, with 10X objective, 1024×1024 resolution, zoom 1, speed 5 fps, line average 4 and Z step 1 µm. The resulting images were produced using ImageJ software.

Brains for cell counting were imaged as follows: Anti-Repo and SarmNP0257> hisYFP were imaged using a Zeiss LSM 710 confocal microscope with a 25x oil immersion lens, at a resolution of 1024×1024 pixels, zoom 0.6 or 1, speed of 7 frames per second (fps), line average of 1 and 0.96 µm step. MyD88>hisYFP samples were imaged using a Leica SP8 confocal microscope with a 20X oil immersion lens, at 1024×1024, zoom 1, speed 400 Hz and Z step 0.96 µm.

Anti-TH+ stained Toll->HisYFP adult brains were imaged with the Leica SP8 confocal microscope with a 20X oil immersion lens, at 1024×1024 pixels, zoom 1.4, speed 400 Hz and Z step 0.96 µm. For anti-TH in wild-type brains, the Zeiss LSM 710 confocal microscope was used, with a 25x oil immersion lens, 1024×1024 at 8 fps, line average of 1 and Z step 0.96 µm. For MyD88>hisYFP anti-TH samples, adult brains were scanned with the Leica SP8 confocal microscope with a 20X oil immersion lens, at 1024×1024, zoom 1.4, speed 400 Hz and Z step 0.96 µm.

### Quantification of cell number using DeadEasy

Cells were counted automatically using DeadEasy software, developed as ImageJ plug-ins, as reported in (Li et al., 2020). Repo+ cells in the adult brain were counted automatically with the *DeadEasy Glia Adult*; Sarm>HisYFP+ and MyD88>HisYFP+ cells were quantified with *DeadEasy adult central brain*; and THGAL4 R58E02GAL4>HisYFP+ cells were counted with *DeadEasy Kenyon Cells*.

### Statistical analysis

Data were collected using *Excel* (Microsoft) and analysed using GraphPad Prism®. Longevity assay data were analysed using Log Rank (Mantel-Cox test). Categorical proboscis extension response data were analysed using Chi-square test. Cell number counting data were numerical and continuous. Datasets with four or more groups were first tested for normality. For sample groups with two sample types, unpaired Student t-tests were conducted; and Mann-Whitney U tests if not normally distributed. For comparisons involving infection vs, not-infected, and two genotypes, Two-Way ANOVA followed by Turkey’s multiple comparison tests were used, using 95% confidence.

## ACKNOWLEDGEMENTS

We are very grateful to Jean Marc Reichhart for advice in the early phases of the project, and to him and Jean-Luc Imler for the gift of *B. bassiana* strains. We thank Guiyi Li, Anna Parsons, Francisca Rojo-Cortés and Xiaocui Wang for constructive criticisms on the manuscript. This work was funded by a Darwin Trust Doctoral Studentship to D.S., BBSRC BB/R017034/1 and Wellcome Trust Investigator Award 223197/Z/21/Z to A.H.

## AUTHOR CONTRIBUTIONS

D.S. carried out all the experiments and collected all the data shown in the figures, was involved in experimental conception and design, data acquisition, curation and analysis. A.R.E.R. and E.Q.M. generated tools. D.A. generated preliminary data. E.B. and H-J.T supervised D.S.; A.H. supervised D.S., A.R.E.R. and D.A., conceived the project and analysed data. D.S., A.H., E.B., H-J.T were involved in experimental design. D.S. and A.H. wrote the manuscript, and all authors contributed to improving the manuscript.

---

**Supplementary Figure 1.**
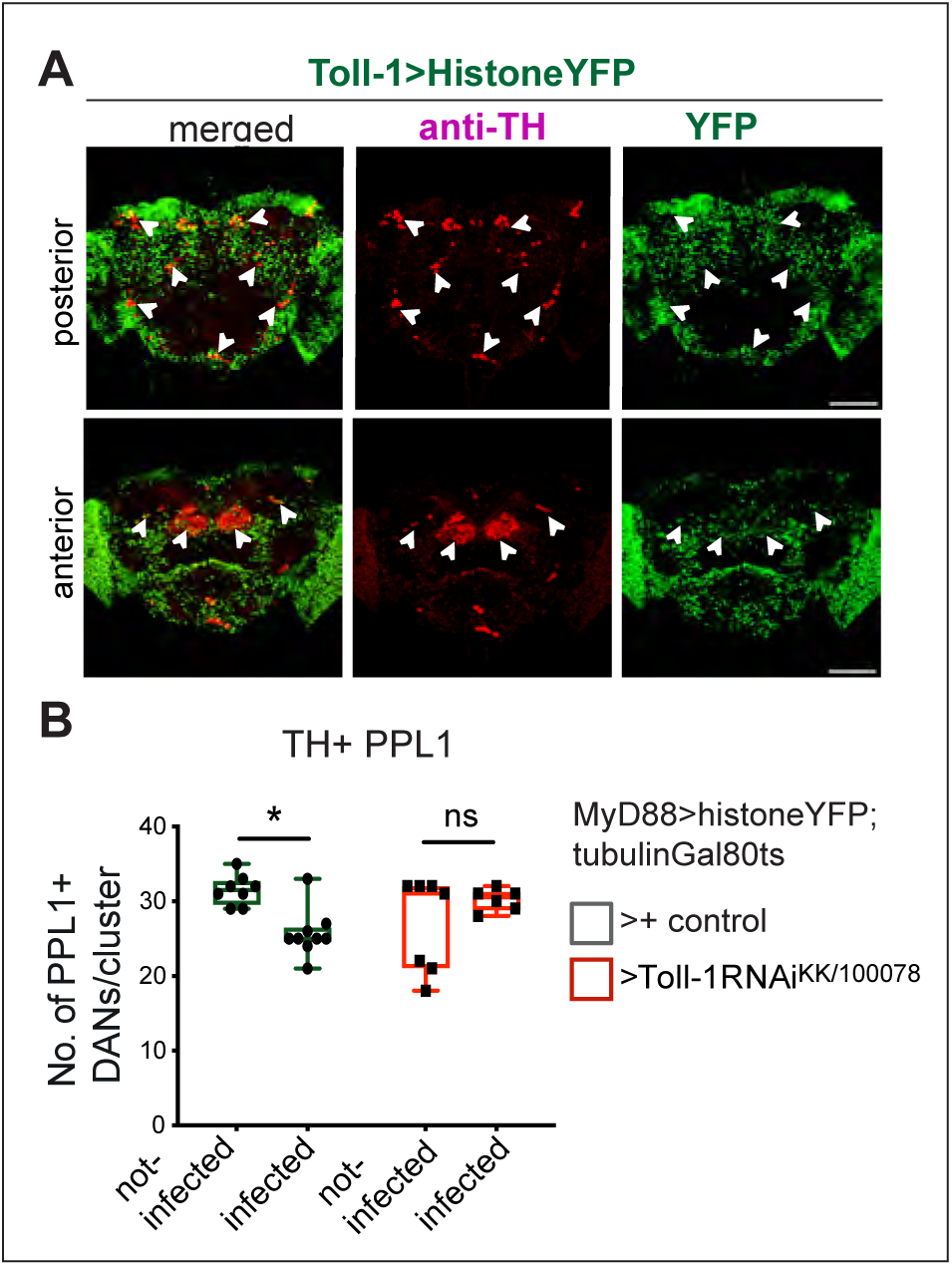
Toll-1 is expressed in PAMs and is required in PPL1. **(A)** Co-localisation of Toll-1>histone-YFP with anti-TH in PPL1, PPL2, PPM3, PAL and a subset of PAM DANs, in adult fly brains. **(B)** Adult-restricted *Toll-1* knock-down in MyD88+ cells does not affect PPL1 neurons, but it rescues the loss of PPL1s caused by *B. bassiana* infection. Two-way ANOVA: Infected vs not-infected p= 0.9586; Control vs UASToll-1RNAi^KK/100078^ (MyD88hisYFP>oregon vs MyD88hisYFP>UASToll-1RNAi^KK/100078^): p= 0.3628; Interaction: p= 0.0024, followed by Turkey’s multiple comparisons correction test. Sample size Toll-1>histone-YFP fly brains n = 5, Oregon>histone-YFP, n = 5, non-infected control brains n = 8, infected control brains n = 9, non-infect*ed Toll-1* ^KK/100078^ *RNAi* brains n= 7, infected *Toll-1* ^KK/100078^ *RNAi* brains n= 6. Asterisks on graphs: *p<0.05, **p<0.01, ***p<0.001, ****p< 0.0001.

**Supplementary Figure 2.**
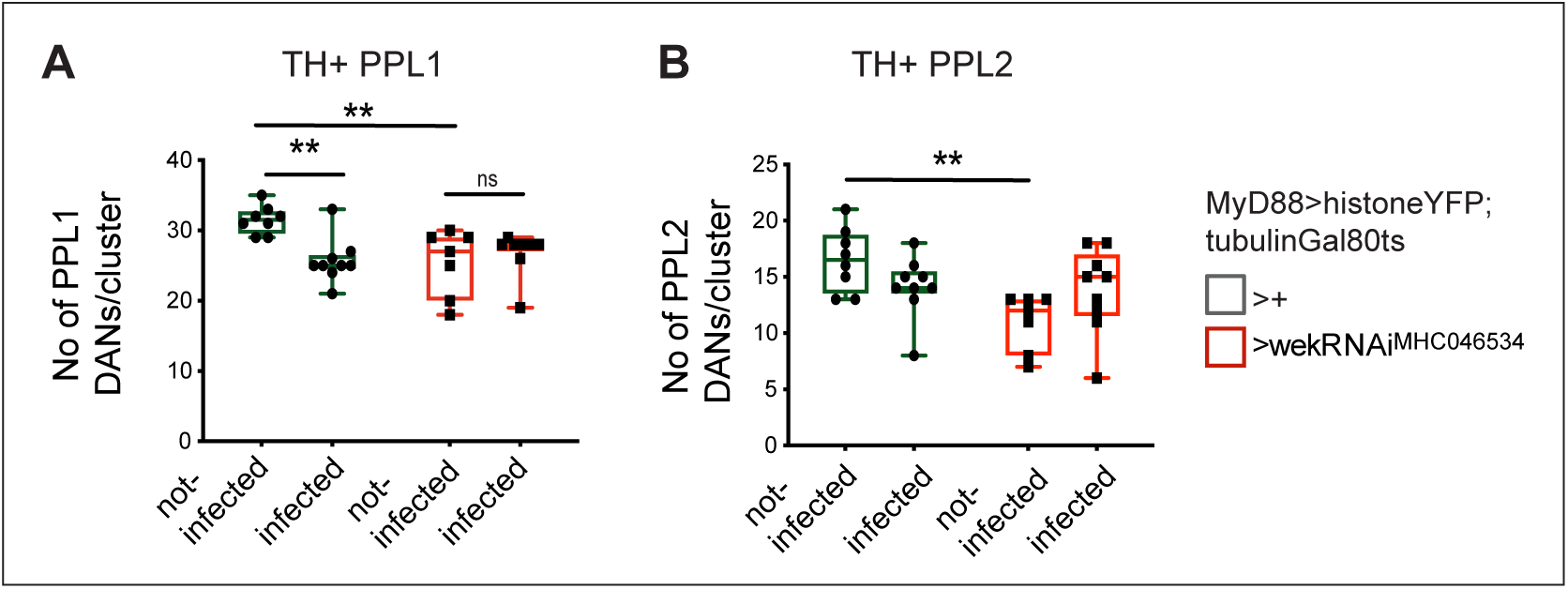
wek knock-down caused DAN loss in the absence of infection. Wek-RNAi knock-down caused a decrease in TH+ DANs PPL1 (A) and PPL2 (B), that was not rescued with the over-expression of wek caused by *B. bassiana* infection, suggesting that wek is required for DAN differentiation. **(A)** Two-way ANOVA: Infected vs not-infected p= 0.0455; Control vs *UASwekRNAi* (*MyD88hisYFP>oregon* vs *MyD88hisYFP>UASwekRNAi*): p= 0.0695; Interaction: p= 0.0038 followed by Turkey’s multiple comparisons correction test: non-infected control brains n = 8, infected control brains n= 9,, non-infected *wek-RNAi* brains n = 7,, infected *wek-RNAi* brains n= 9. (B) Two-way ANOVA: Infected vs not-infected p= 0.0104; Control vs *UASwekRNAi (MyD88hisYFP>oregon vs MyD88hisYFP> UASwekRNAi)*: p= 0.8564; Interaction: p= 0.0217 followed by Turkey’s multiple comparisons correction test: non-infected control brains n = 8, infected control brains n= 9, non-infected *wek-RNAi* brains n = 7,, infected *wek-RNAi* brains n= 9. Asterisks on graphs: *p<0.05, **p<0.01, ***p<0.001, ****p< 0.0001.

## Notes

### Competing Interest Statement

The authors have declared no competing interest.

